# A broad-specificity O-glycoprotease that enables improved analysis of glycoproteins and glycopeptides containing intact complex O-glycans

**DOI:** 10.1101/2021.09.09.459412

**Authors:** Saulius Vainauskas, Hélène Guntz, Elizabeth McLeod, Colleen McClung, Cristian Ruse, Xiaofeng Shi, Christopher H. Taron

## Abstract

Analysis of mucin type O-glycans linked to serine/threonine of glycoproteins is technically challenging, in part, due to a lack of effective enzymatic tools that enable their analysis. Recently, several O-glycan-specific endoproteases that can cleave the protein adjacent to the appended glycan have been described. Despite significant progress in understanding the biochemistry of these enzymes, known O-glycoproteases have specificity constrains, such as inefficient cleavage of glycoproteins bearing sialylated O-glycans, high selectivity for certain type of glycoproteins or protein sequence bias, that limit their analytical application. In this study, we examined the capabilities of an immunomodulating metalloprotease (IMPa) from *Pseudomonas aeruginosa*. The peptide substrate sequence selectivity and its impact on IMPa activity was interrogated using an array of synthetic peptides and their glycoforms. We show that IMPa has no specific P1 residue preference and can tolerate most amino acids at the P1 position, except aspartic acid. The enzyme does not cleave between two adjacent O-glycosites, indicating that O-glycosylated serine/threonine is not allowed at position P1. Glycopeptides with as few as two amino acids on either side of an O-glycosite were specifically cleaved by IMPa. Finally, IMPa efficiently cleaved peptides and proteins carrying sialylated and asialylated O-glycans of varying complexity. We present the use of IMPa in a one-step O-glycoproteomics workflow for glycoprofiling of individual purified glycoproteins granulocyte colony-stimulating factor (G-CSF) and receptor-type tyrosine-protein phosphatase C (CD45) without the need for glycopeptide enrichment. In these examples, IMPa enabled identification of O-glycosites and the range of complex O-glycan structures at each site.

## Introduction

Glycosylation of proteins is a common and complex post-translational modification that plays important roles in numerous biological processes.^1–3^ Over one hundred glycosyltransferases and other modifying enzymes are involved in synthesis of different types of mammalian glycans via multiple biosynthetic pathways. The combinatorial action of these enzymes on conserved protein and lipid scaffolds creates both micro- and macroheterogeneity of glycosylated molecules within the mammalian glycome.

N-Glycosylation and mucin-type O-glycosylation of proteins are the two major classes of carbohydrate modification found on eukaryotic extracellular membrane and secretory proteins. Glycan structures and their site of glycan attachment in a polypeptide (a ‘glycosite’) significantly differ between these two classes of glycosylation.^2–3^ N-Glycan precursors are assembled and transferred to an asparagine sidechain within the canonical consensus sequence N-X-S/T in the endoplasmic reticulum. Protein-bound N-linked glycans are further modified by Golgi-localized glycosyltransferases. O-Glycosylation occurs in the Golgi, where it is initiated by addition of N-acetylgalactosamine (GalNAc) to a side-chain hydroxyl of serine or threonine residues in a protein. There is no clear protein consensus sequence that defines O-glycosites, making their identification challenging. Furthermore, the absence of a robust enzyme that can release intact complex O-glycans further complicates characterization of O-glycosylation.

To overcome these technical hurdles, a new glycoproteomic approach to O-glycan and O-glycosite analyses has been emerging. Central to this method is the use of bacterial O-glycan specific proteases (O-glycopeptidases or O-glycoproteases).^4–8^ O-glycoproteases are a growing group of highly specific enzymes that recognize and cleave a glycopeptide/glycoprotein adjacent to a O-glycosylated serine or threonine. Some O-glycopeptidases hydrolyze the peptide bond exclusively N-terminal to an O-glycosylated Ser/Thr residue, whereas others cleave on the C-terminal side.^5,9–11^ MS analysis of glycopeptides generated by O-glycoprotease digestion provides important information about the location of O-glycosites within a protein and the structure of O-glycans present at each O-glycosite. Despite significant progress in understanding this new class of enzymes, known O-glycopeptidases have applied limitations. For instance, the O-glycopeptidases OgpA from *Akkermansia municiphila* and BT4244 from *Bacteroides thetaiotaomicron* have limited or no ability to hydrolyze sialylated glycopeptides.^6,9–10^ A mucin-specific protease StcE from an enterohemorrhagic *Escherichia coli* strain selectively recognizes and cleaves heavily glycosylated mucin domains but shows no activity on non-mucin O-glycoproteins.^5^ The activity of the glycoprotease CpaA from *Acinetobacter* species is not affected by the presence of sialic acids, but it hydrolyzes the peptide backbone only next to O-glycosites preceded by proline (Pro-Ser/Thr).^11^ A zinc metalloprotease C (ZmpC) from *Streptococcus pneumoniae* is also not affected by sialic acids but is selective for mucins and cleaves the peptide backbone several amino acids away from an O-glycosylated serine or threonine residue.^7^ In all cases, valuable information about the location of O-glycosites and/or structure of O-glycans present at each glycosite may be lost using any of the aforementioned O-glycoproteases in an analytical workflow, providing an incomplete picture of a protein’s O-glycosylation content.

In this study, we examined the capabilities of a different bacterial enzyme, an immunomodulating metalloprotease from *Pseudomonas aeruginosa* (IMPa).^12^ The enzyme cleaves immediately N-terminal to a O-glycosylated serine or threonine residue. It has been previously reported that this O-glycoprotease is able to recognize a variety of O-glycan structures and that its activity is not affected by the presence of sialic acids on the glycan.^9–10^ It has been recently proposed that the N-terminal portion of IMPa contains a proline-specific recognition domain that may target the protease to proteins containing an O-glycosylated Pro-Ser/Thr motif.^13^ Crystallographic analysis of the native IMPa with glycosylated serine residue revealed the specific interactions between glycan and amino acids in the active site^9^, however, the peptide substrate sequence selectivity and its impact on IMPa activity need further elucidation.

In this work, we undertook biochemical characterization of IMPa specificity using an array of synthetic glycopeptides and digestion of O-glycosylated proteins. We investigated the importance of each amino acid at the P1 position of the substrate peptide for the IMPa’s activity. We show that IMPa acts as a broad-specificity O-glycoprotease with minimal influence of amino acid composition adjacent to the cleavage site. Finally, using purified glycoproteins with varying levels of O-glycosylation, we show that IMPa effectively generates O-glycopeptides that can be used to profile O-glycosites and determine the range of glycans present at each position. This work helps to better understand IMPa function and illustrates its potential in analysis of individual O-glycoproteins, including biotherapeutics.

## Experimental Procedures

### Materials

All chemical reagents and solvents were from MilliporeSigma (Burlington, MA) unless otherwise stated. RapiFluor-MS was from Waters (Milford, MA). Etanercept (Enbrel^®^) was obtained from Myoderm (Norristown, PA), recombinant human CD45 (cat no. 1430-CD-050) was from R&D Systems (Minneapolis, MN), recombinant human CD45 (cat no. 14197-H08H) was from SinoBiological (Wayne, PA), recombinant human G-CSF (cat no. CYT-088) was from ProSpec (East Brunswick, NJ). O-Glycoprotease (IMPa from *Pseudomonas aeruginosa*), Trypsin-Ultra, O-Glycosidase, α2-3,6,8 Neuraminidase, and α2-3,6,8,9 Neuraminidase A were all from New England Biolabs (Ipswich, MA). OpeRATOR (OgpA from *Akkermansia municiphila*) was from Genovis Inc (Cambridge, MA). Fetuin from fetal bovine serum was obtained from Sigma-Aldrich.

### Preparation of O-glycopeptides for capillary electrophoresis (CE) analysis

A fluorescent glycopeptide (AHGVT*SAPDTRK-FAM) of 12 amino-acids with an O-glycan tri-saccharide (Neu5Ac-α2,3-Gal-β1,3-GalNAc) on the first Thr residue (*), and a C-terminal fluorescein amidite group (FAM) was obtained through custom synthesis (Bio-Synthesis Inc., Lewisville, TX). To generate a glycopeptide substrate containing asialylated TF-antigen (Gal-β1,3-GalNAc), the synthetic O-glycopeptide was incubated with 200 U of α2-3,6,8 Neuraminidase (New England Biolabs) for 16 h at 37°C. To prepare a glycopeptide bearing a T-antigen (GalNAc) glycan, the glycopeptide was incubated with 200 U of α2-3,6,8 Neuraminidase (New England Biolabs) and 40 U of β1-3 Galactosidase (New England Biolabs) for 16 h at 37°C.

### O-glycoprotease activity assay

For capillary electrophoresis (CE) analysis, 80 pmol of FAM-labeled glycopeptide substrate was mixed with 1 μg of O-glycan specific protease in 10 μl of 20 mM Tris-HCl, pH 8.0 and incubated for 5 h at 37°C. Following incubation, the reaction products were diluted 200-fold with water and analyzed by CE.

### Capillary electrophoresis (CE)

Glycopeptide samples were separated using an 3730xl DNA Analyzer with a 96 capillary (36 cm) array (Applied Biosystems, Waltham, MA) filled with Performance Optimized Polymer (POP7, Thermo Fischer Scientific, Waltham, MA). The oven temperature was 60°C, the buffer temperature was 35°C, and the current stability was 30 µA. A pre-run was done at 15 kV for 180 s. Sample injection was done at 3 kV for 15 s after which, the sample was separated at 15 kV for 1800 s. Data were collected by a charge-coupled device (CCD) camera and analyzed with the Peak Scanner software version 1.0 (Thermo Fischer Scientific).

### Proteolytic digestion

Native glycoproteins (10 μg) were digested with 1 U of IMPa (O-Glycoprotease, New England Biolabs) in 50 μL 20 mM Tris-HCl, pH 8.0 for 5 h at 37°C. After incubation, glycoprotein hydrolysates were passed through a Microcon-30kDa centrifugal filter unit (MilliporeSigma, Burlington, MA) and dried in a SpeedVac (Thermo Fisher Scientific). Dried peptides were resuspended in water and analyzed by LC-MS/MS or labeled with RapiFluor-MS (RFMS) or 6-aminoquinolyl-N-hydroxysuccinimidyl carbamate (AQC).

### Peptide derivatization with RapiFluor-MS, AQC

A RapiFluor-MS (Waters, Milford, MA) stock solution was prepared by dissolving 9 mg of RapiFluor-MS in 130 μl of dimethylformamide. A 21 μl aliquot of this stock was added to peptides (5-10 μg) that had been resuspended in 10 μl of 25 mM sodium bicarbonate. The reaction was incubated for 5 min at room temperature in the dark. Labeled peptides were cleaned through a hydrophilic interaction chromatography solid phase extraction (HILIC SPE) MacroSpin column (Nest Group Inc., Southborough, MA). Briefly, labeling samples were diluted to 300 μL with 90% acetonitrile/10% 50 mM ammonium formate, pH 4.4 (90% ACN/CH_5_NO_2_), loaded on columns equilibrated with the same buffer, and washed 5 times with 350 μL 90% ACN/CH_5_NO_2_. Samples were eluted in 100 μL of 50 mM ammonium formate, pH 4.4 and dried in a SpeedVac (Thermo Fisher Scientific).

A 10 mM 6-aminoquinolyl-N-hydroxysuccinimidyl carbamate (AQC) solution was prepared by dissolving 3 mg of AQC in 1 mL of acetonitrile. A 24 μL aliquot of 10 mM AQC was added to the peptides (5-20 μg) that had been resuspended in 10 μL of 25 mM sodium bicarbonate. The mixture was incubated for 10 min at 55°C in the dark. Labeled peptides were cleaned through HILIC SPE MacroSpin columns (Nest Group Inc.) as described above.

### UPLC-HILIC-FLR-MS1

RapiFluor-MS-labeled or AQC-labeled samples were separated by UPLC using a ACQUITY UPLC glycan BEH amide columns 130Å (2.1 × 150 mm, 1.7 μm) or 300Å (2.1 × 150 mm, 1.7 μm) from Waters on a H-Class ACQUITY instrument (Waters, Milford, MA). Solvent A was 50 mM ammonium formate buffer, pH 4.4 and solvent B was acetonitrile. The gradient used with 130Å (2.1 × 150 mm, 1.7 μm) column was 0–40 min, 10–46% solvent A; 41.5–44.5 min, 100% solvent A; 48.1–55 min, 10% solvent A. The flow rate was 0.4 mL/min. The gradient used with 300Å (2.1 × 150 mm, 1.7 μm) column was 0–35 min, 15–50% solvent A; 35–36 min, 100% solvent A; 36–38 min, 100%; 38–39 min, 15%; 39–49 min, 15% solvent A. The flow rate was 0.2 mL/min. The injection volume was 18 μL and the sample was prepared in 90% (v/v) acetonitrile. Samples were kept at 5°C prior to injection and the separation temperature was 60°C. The fluorescence detection wavelengths were λ_ex_= 265 nm and λ_em_= 425 nm for RapiFluorMS and λ_ex_= 246 nm and λ_em_=396 nm for AQC, with a data collection rate of 20 Hz.

Conditions for inline mass detection using the ACQUITY quadrupole QDa (Waters) were as follows: Electrospray ionization (ESI) in positive mode; capillary voltage, 1.5 kV; cone voltage, 15 V; sampling frequency, 5 Hz; probe temperature 400°C. The QDa analysis was performed using full scan mode, and the mass range was set at m/z 100–1250. Single ion recording (SIR) mode was used as well to monitor individual glycopeptides. Waters Empower 3 chromatography workstation software was used for data processing including traditional integration algorithm, no smoothing of the spectra and manual peak picking.

### Nano LC-MS/MS analysis

Samples were directly injected on a reversed phase 1.9 μm C18 fused silica column (I.D. 75 μm, O.D. 360 μm × 25 cm length) packed into an emitter tip (IonOpticks, Australia). The column was placed in a column oven integrated into the mass spectrometer ion source (Sonation Column Oven). The flow rate for analysis was 400 nL/min with buffer A (water, 0.1% formic acid) and buffer B (acetonitrile, 0.1% formic acid). The gradient conditions were: 2 min at 2% B, 30 min to 40% B, 5 min to 70% B, 2 min at 70% B with 1 min return to 2% B and a total of 40 min analysis time. The temperature was set at 40°C. The Easy-nLC 1000 was coupled with a QExactive Hybrid Quadrupole-Orbitrap mass spectrometer. The most abundant 10 ions were fragmented in the normalized collision energy NCE 27.

### Data analysis

Glycopeptide MS/MS fragment spectra were searched against selected protein and glycan databases using Byonic software v4.0-53 x64 (Protein Metrics Inc.).^14^ Searches were performed using precursor ion mass tolerance of 6 ppm, fragment ion mass tolerance of 30 ppm, specific or semi-specific cleavage at the N-terminus of Ser/Thr with up to 7 missed cleavages allowed. Variable modifications were used in searches: oxidation of Met (+15.994 Da), deamidation at Asn, Gln (+0.984 Da). An O-glycan structure database containing 13 common mammalian O-glycans was used (Table S1). Up to two glycan modifications were allowed per peptide. Searches were performed against defined glycoproteins to narrow the search space. The searches were filtered using strict criteria (1% FDR, PEP 2D < 0.01). Validation of the identified glycopeptides using oxonium ions of *m/z* 204.087 (HexNAc), *m/z* 284.044 (sulfoHexNAc), *m/z* 274.092/292.103 (Neu5Ac), *m/z* 316.103/334.114 (acetylNeu5Ac), *m/z* 290.087/308.098 (Neu5Gc), *m/z* 407.166 (2HexNAc), *m/z* 569.219 (2HexNAcHex) in the MS^2^ spectrum was performed to ensure correct assignment of O-glycopeptides.

## Results

### Specificity comparison of different O-glycoproteases

The specificities of different O-glycan-specific proteases were compared using a synthetic peptide derived from the MUC1 protein that carries different core 1 O-glycan forms. Proteases with previously reported selectivity for cleavage of different O-glycoproteins or O-glycopeptides were chosen for comparison with an immunomodulating protease (IMPa, supplied commercially as ‘O-Glycoprotease’) from *Pseudomonas aeruginosa*.^9,12^ BT4244 from *Bacteroides thetaiotaomicron* was previously shown to have proteolytic activity on O-glycosylated peptides and proteins.^9,15^ An O-endoprotease OgpA from *Akkermansia muciniphila* (supplied commercially as ‘OpeRATOR’) has been used for digestion of mucin-type glycoproteins and glycopeptides.^4,6^ Each of these proteases cleave the peptide bond immediately N-terminal to the serine or threonine residue that harbors an O-glycan.^4,9,15^ A fourth enzyme tested was a mucin-specific bacterial protease StcE (secreted protease of C1-esterase inhibitor) from *Escherichia coli*.^16^ It was previously reported that StcE recognizes and cleaves mucin-type glycoproteins on the C-terminal side of serine or threonine residues that possess an O-glycan.^5^

To evaluate activity of these enzymes, fluorescent O-glycopeptide substrates were incubated with recombinantly produced and purified enzymes (Supporting Information S1 and Figure S1). Reaction products were analyzed using capillary electrophoresis (CE) (Figure S2). Only IMPa efficiently cleaved all three glycopeptides containing different O-glycans (97-99% efficiency), whereas other tested O-glycopeptidases digested the glycopeptides with varying degrees of efficacy (Table 1). BT4244 from *Bacteroides* cleaved glycopeptides containing TF- and T-antigen but exhibited no activity on peptides possessing sialylated TF-antigen, whereas OgpA from *Akkermansia* efficiently digested asialylated TF-antigen carrying peptide but showed little activity on glycopeptide with sialylated TF-antigen and no activity on a glycopeptide containing T-antigen (Table 1). No digestion of the mucin glycopeptides by StcE was detected in this assay. Additionally, none of these proteases were active on a control peptide that lacks an O-glycan, emphasizing that in this assay, observed activity of IMPa, BT4244, and OgpA is dependent on the presence of an O-glycan (Figure S3). However, because IMPa was the only enzyme that showed the ability to efficiently cleave each glycopeptide we tested regardless of the complexity of the appended O-glycan, we focused our efforts on further defining the specificity of this protease.

**Table 1.**
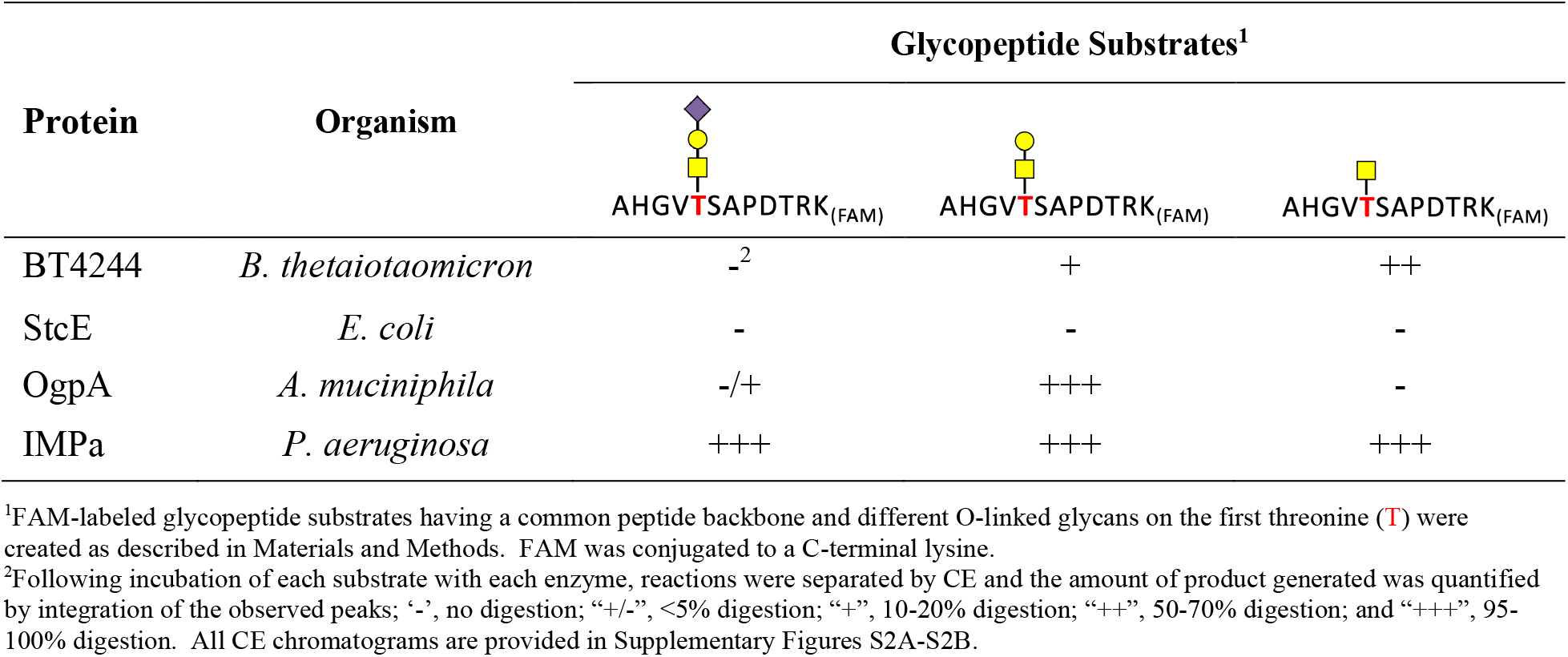
Activity of known O-proteases on FAM-labeled glycopeptide substrates.

### IMPa digestion of glycopeptide variants with different amino acids at the P1 position

To investigate if IMPa cleavage is affected by a specific amino acid residue at the position P1 of O-glycopeptide substrates, its activity on a panel of synthetic peptides containing a different residue at the position P1 was examined. An 11 aa peptide (PPDAASAAPLR) derived from erythropoietin, a protein known to contain a single O-glycosite, was used as a scaffold to create a family of related enzyme substrates. Twenty peptides that differed only in a single amino acid at P1, were synthesized and derivatized with the fluorescent label 6-aminoquinolyl-N-hydroxysuccinimidyl carbamate (AQC). This compound utilizes reactive NHS-carbamate chemistry to form a covalent urea linkage between the fluorophore and free amino groups and can efficiently modify a peptide at its N-terminus for fluorescence detection during UPLC-HILIC-FLR analysis (Figure 1C). AQC-labeled peptides were each incubated with yeast expressed recombinant human polypeptide N-acetylgalactosaminyltransferase 1 (GALNT1) (Supporting Information S2 and Figure S4) in the presence of UDP-GalNAc to generate O-glycopeptides containing O-linked GalNAc (Tn-antigen) on the serine residue (Figure 1). Each glycopeptide and its corresponding aglycosylated counterpart were incubated overnight with IMPa. The reaction products were analyzed by UPLC-HILIC-FLR-MS (Figure 1C,D and Figures S5-S15). The fluorescent label at the N-terminus of each peptide permitted detection and relative quantification of the glycopeptide substrate, and the peptide product generated upon endoproteolytic digestion (Table 2). The identity of peptide products was confirmed using a Quadrupole Dalton (QDa) mass detector coupled with in-line fluorescence detection (UPLC-HILIC-FLR-MS) (Figure 1D).

**Table 2.** P1 residue specificity of IMPa.

| P1<br>amino acid | IMPa activity<br>% |
| --- | --- |
| A | 100 |
| V | 96 |
| Y | 90 |
| W | 79 |
| T | 100 |
| P | 97 |
| F | 89 |
| L | 92 |
| I | 8 |
| H | 83 |
| G | 100 |
| E | 98 |
| C | 100 |
| D | nd |
| N | 98 |
| R | 38 |
| M | 100 |
| K | 100 |
| S | 100 |
| Q | 94 |

nd - not detected

**Figure 1.**
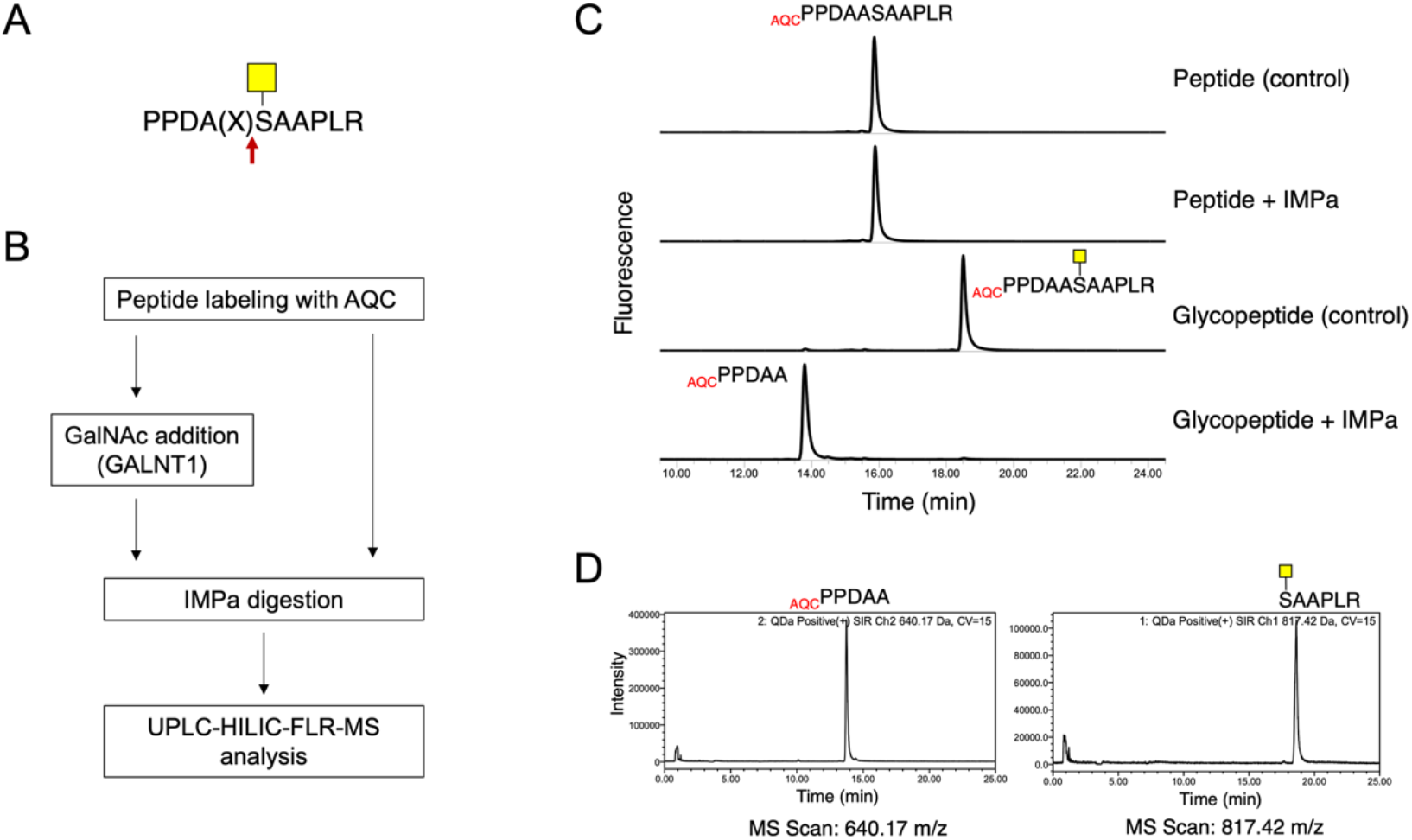
Approach for generation and analysis of O-glycopeptide enzyme substrates. (A) A set of 20 O-glycopeptides having the core sequence PPDA(X)SAAPLR was constructed. Symbols: (X), the P1 position where each of the 20 amino acids was inserted; red arrow, IMPa cleavage site; yellow box, α-GalNAc. (B) A schematic depicting the experimental process of glycopeptide labeling, IMPa digestion, and peptide analysis. (C) An example UPLC-HILIC-FLR chromatogram showing separation of the PPDAASAAPLR peptide (control) and its corresponding O-glycopeptide before and after IMPa digestion. (D) Analysis of the glycopeptide reaction products from panel C (bottom chromatogram) by UPLC-HILIC with inline MS after digestion with IMPa. The reaction product (GalNAc)SAAPLR is detected by MS but not by UPLC-HILIC-FLR (panel C) due to removal of the AQC label upon IMPa digestion.

Endoproteolytic cleavage at the N-terminus of Ser was observed solely with O-glycopeptide substrates, but not with non-glycosylated control peptides (Figure 1C and Figures S5-S15). Most glycopeptides were cleaved with high efficiency (83-100% completion). However, cleavage of glycopeptides containing Arg or Ile in the P1 position was less effective (38% and 8% completion, respectively). The glycopeptide containing Asp at P1 was the only member of this set not cleaved by IMPa (Table 2). This observation is also supported by data from a prior study where IMPa did not digest a different O-glycopeptide containing Asp at P1 position and an O-glycan on Thr at P1’.^9^ Thus, it is likely that IMPa does not cleave O-glycopeptides when Asp is in the P1 position, followed by glycosylated Ser or Thr (P1’ position).

### Recognition of different glycopeptide substrates by IMPa

Extended incubation (18-24 h) of a glycopeptide having threonine in the P1 position (PPDATSAAPLR) with GALNT1 resulted in GalNAc modification of both adjacent threonine and serine residues (Figure 2A and Figure S14B). Shorter incubation (4 h) of the same substrate with GALNT1 produced a heterogeneous mixture of two peptides, one having a single GalNAc at Ser and a second having GalNAc at both Thr and Ser (Figure S14A). Both of these glycopeptides were efficiently cleaved by IMPa. Interestingly, the glycopeptide with two adjacent O-glycans was cleaved only on the N-terminal side of Thr (Figure 2A and Figure S14). No reaction products indicating cleavage between O-glycosylated Thr and Ser were observed even after extended 24 hours incubation with IMPa, suggesting that IMPa does not cut between two adjacent O-glycosites.

**Figure 2.**
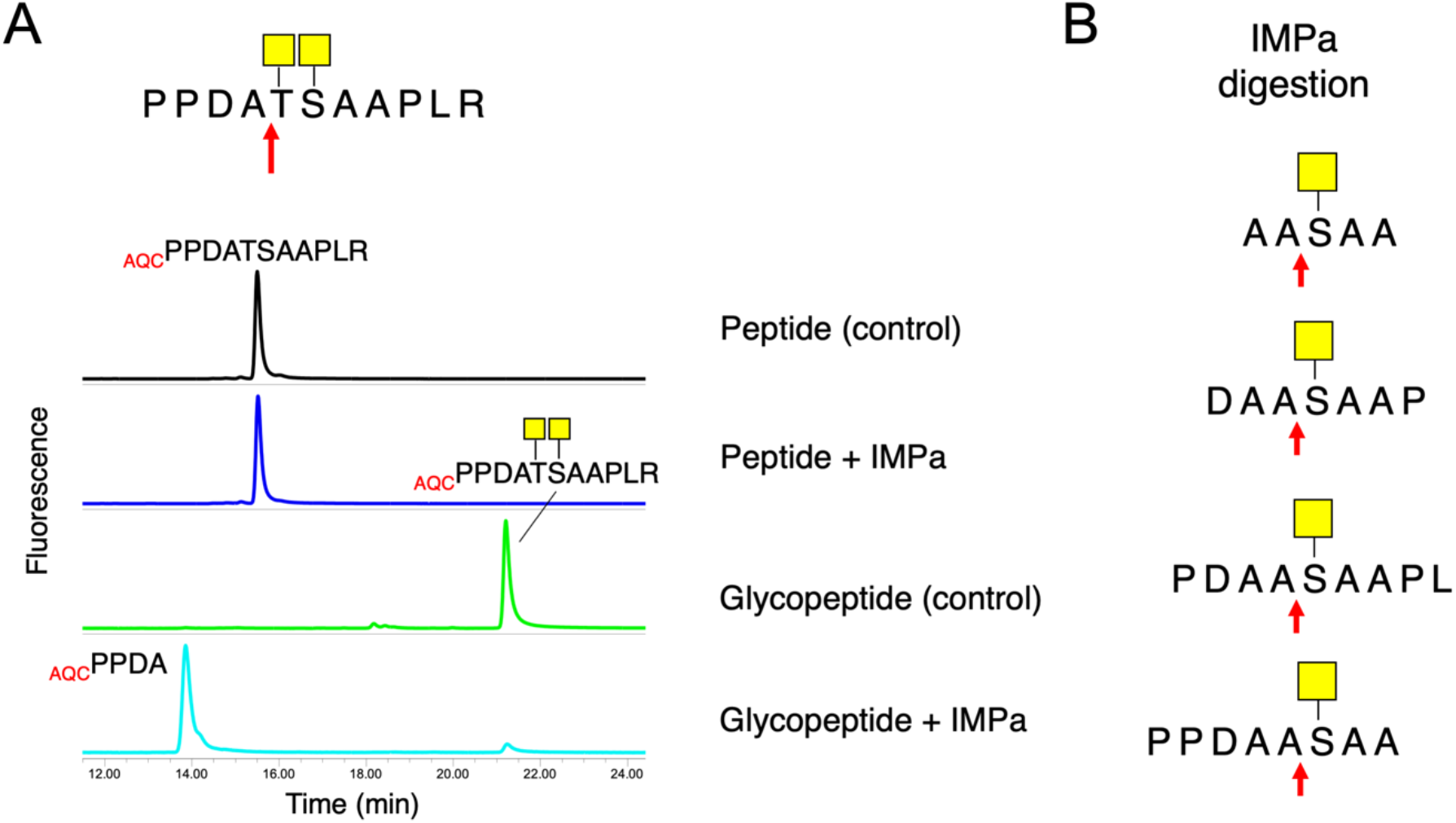
Peptide substrate specificity of IMPa. (A) Observed IMPa cleavage of a glycopeptide containing two adjacent O-glycosites. (B) Structures of nested small glycopeptide substrates that were cleaved by IMPa (see Figures S16-S18 for more information). Each substrate contains progressively fewer amino acid residues on the amino terminal side of the O-glycosite. The arrow depicts observed IMPa cleavage site.

The minimum length of a peptide required for cleavage by O-glycoprotease was also investigated using synthesized nested truncations of the erythropoietin peptide. Peptides of 5 (AASAA), 7 (DAASAAP), 8 (PPDAASAA) and 9 (PDAASAAPL) amino acids with each having 2-4 amino acids on either the N-terminal or C-terminal side of the O-glycosite (Ser) were synthesized, AQC-labeled and glycosylated *in vitro* as described above (Figure 1B). GalNAc addition to two peptides (AASAA and PPDAASAA) was inefficient with only <5% transfer occurring. This suggests that these two peptides are poor substrates for GALNT1 *in vitro*. All glycopeptides (including the two with only partial GalNAc addition) were incubated with IMPa. Specific cleavage at the N-terminus of O-glycosylated serine was detected for all five tested glycopeptides indicating that as few as two amino acids on either side of an O-glycosite is sufficient for catalysis (Figure 2B and Figures S16-S18). No digestion was detected for any non-glycosylated control peptides (Figure S16-S18).

### Digestion of intact glycoproteins by IMPa

The *in vitro* glycopeptide specificity of IMPa was compelling and its suitability for use in glycoproteomic workflows was further investigated. For IMPa to be suitable for use in such analyses, it would need to be able to generate glycopeptides from larger protein substrates. Thus, several model glycoproteins and recombinant biotherapeutic proteins were used to assess the ability of IMPa to generate glycopeptides from intact glycoproteins. Granulocyte colony-stimulating factor (G-CSF), receptor-type tyrosine-protein phosphatase c (CD45), bovine fetuin and etanercept (ENBREL) are each known to contain a varying number of highly sialylated O-glycans.^17–22^ For instance, over a dozen O-glycosites have been identified on etanercept, six on bovine fetuin, and a single site on recombinant G-CSF. The most complex protein was the longest form of human recombinant CD45 (CD45RABC) that has over 60 potential O-glycosylation sites predicted using the NetOGlyc 4.0 server (https://services.healthtech.dtu.dk/service.php?NetOGlyc-4.0).^23^ Only a small fraction of these glycosites have been experimentally verified on various CD45 isoforms.^4,5,24,25^

The four substrate glycoproteins were each incubated with IMPa, and the generated peptides were separated from the protease by passage through a centrifugal filter device. Peptides were labeled with the RapiFluor-MS (RFMS) fluorescent dye and analyzed by UPLC-HILIC-FLR. Distinct chromatographic profiles of digested peptides were detected for each glycoprotein sample after incubation with IMPa (Figure 3). Observed peptide retention time shifts in response to treatment with neuraminidase and O-glycosidase confirms that they are O-glycopeptides (Figure 3). Additionally, for each protein, the number of observed O-glycopeptides (chromatographic peaks) generally correlated with the known or predicted number of O-glycosites in each glycoprotein.

**Figure 3.**
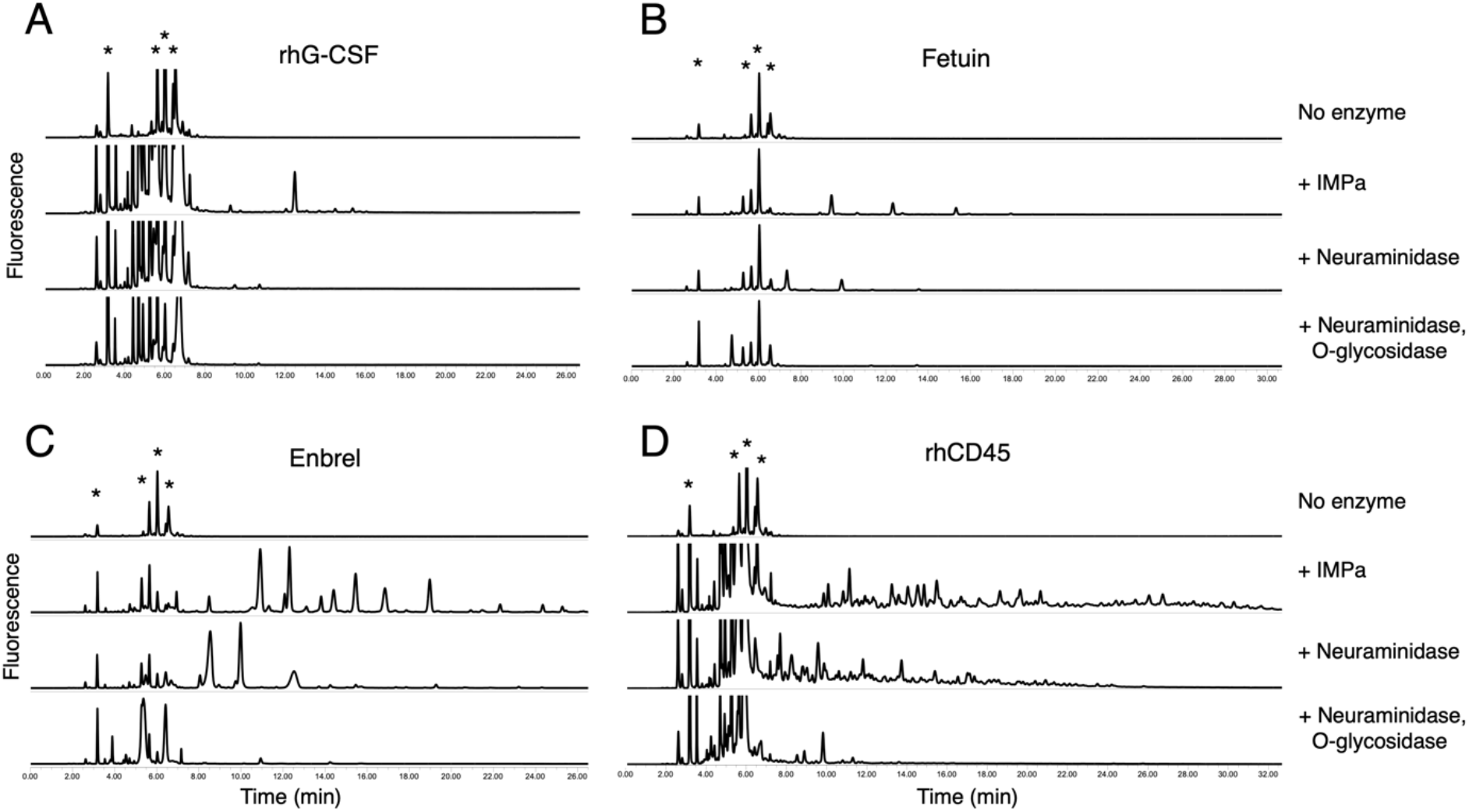
Generation of peptides from intact O-glycoproteins by IMPa digestion. Purified proteins (5-10 µg) A) rhG-CSF, B) bovine fetuin, C) Enbrel, and D) rhCD45 were digested with IMPa (5 h at 37°C). Generated peptides were labeled with RapiFluor-MS and analyzed by UPLC-HILIC-FLR (second chromatogram from the top in each panel), and subsequently incubated with neuraminidase A alone (third chromatogram) or in combination with O-glycosidase (fourth chromatogram). Asterisks denote free RapiFluor-MS and its adducts.

These data support the notion that IMPa treatment of glycoproteins generates an abundance of O-glycopeptides (most carrying sialylated O-glycans), suggesting the enzyme could be used as a first step in a O-glycoproteomics analytical workflow. Thus, we explored this notion using IMPa digestion and nano LC-MS/MS analysis of both the simplest (G-CSF) and most complex (CD45) O-glycoproteins used in the peptide generation experiment (above).

### Site-specific O-glycoprofiling of recombinant human G-CSF (rhG-CSF)

A single O-glycosite has been previously reported on recombinant human G-CSF (rhG-CSF) expressed in CHO cells or in yeast.^17,26^ This glycoprotein stimulates bone marrow production of granulocytes and stem cells and plays essential functions in maintaining the circulating level of active neutrophils.^27,28^ Its recombinant forms (lenograstim or filgrastim) have been used as a therapeutic to treat patients who have had bone marrow transplantation or have developed neutropenia after chemotherapy. The rhG-CSF protein produced in human embryonic kidney (HEK 293) cells was analyzed in this study.

Digestion profile of rhG-CSF with IMPa was analyzed by nano LC-MS/MS. Digested peptide mixtures were directly analyzed by loading onto an analytical reversed phase C18 column coupled to quadrupole-Orbitrap mass spectrometer equipped with a higher energy collisional dissociation (HCD) fragmentation. Glycopeptide MS/MS fragment spectra were searched against selected protein and glycan databases using Byonic software. Our glycan searches used a small database comprised of 13 most common mammalian O-glycans structures that contain Neu5Ac or Neu5Gc (structures are listed in Table S1). Additionally, up to two O-glycan modifications per peptide were permitted to maintain practical search times. The abundance of the detected O-glycan structures was calculated using only high confidence annotations (mostly identified glycopeptides possessing single O-glycan). O-Glycosite and O-glycan structure assignments were considered unambiguous when glycopeptide fragmentation spectra provided sufficient evidence (*e*.*g*., peptide backbone and glycan fragment ions, glycan oxonium ions, fragments with glycans that enable glycosite localization) to support them. In addition, the IMPa cleavage specificity rules (*e*.*g*., N-terminal O-glycosite location on IMPa-generated peptide; no digest between two adjacent O-glycosites) were applied to check each software annotated O-glycosite on the peptide.

A total of 490 peptide spectrum matches (PSM’s) were mapped to G-CSF protein. Of these, 324 PSM’s (66% of total) were assigned to 19 unique glycopeptides and 37 glycoforms of these peptides (Table S2). Three O-glycosites (Thr31, Ser38 and Thr166) were assigned unambiguously. The assignment of a fourth O-glycosite at Thr148 was based on the strict IMPa cleavage specificity, since not enough fragmentation spectral information was obtained for unambiguous assignment of the O-glycosite on the identified glycopeptide (Figure 4 and Table S2). Only a single O-glycosite (Thr166) has been identified previously on rhG-CSF expressed in CHO cells or in yeast.^17,26^ However, our data show that additional O-glycosites are present on rhG-CSF produced by HEK 293 cells.

**Figure 4.**
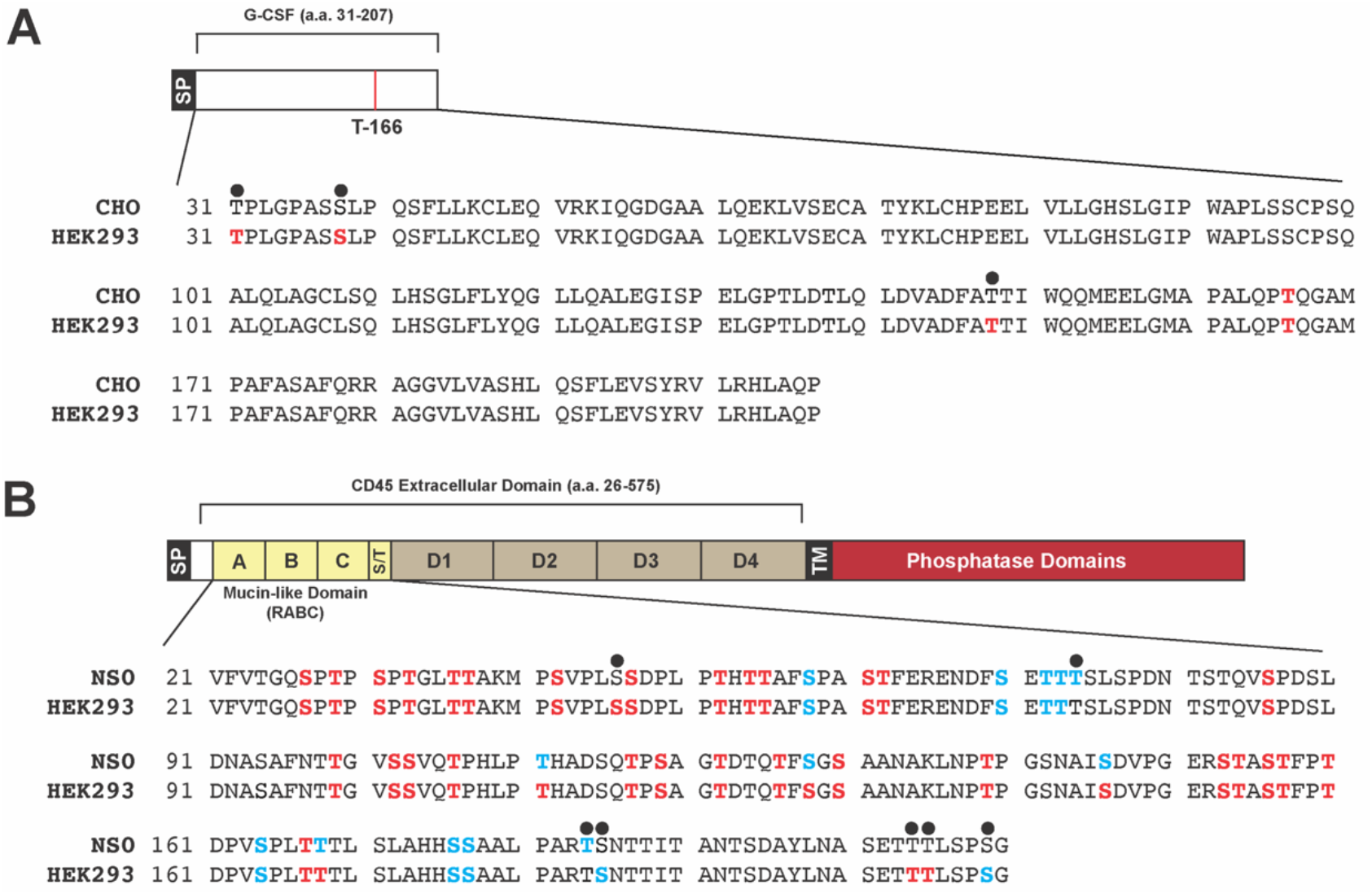
Mapping O-glycosites in rhG-CSF and rhCD45. The recombinant proteins (**A**) rhG-CSF produced in HEK 293 cells and (**B**) rhCD45 produced in NS0 and HEK 293 cell lines, were treated with IMPa and the resulting peptides analyzed by LC-MS/MS as described in Materials and Methods. Shown are the locations of observed O-glycosites (red letters) and “inferred” O-glycosites (blue letters) for each protein. A site was considered inferred when no peptide starting at the prospective site was observed, but a peptide mapping immediately upstream terminated at the prospective site (suggesting IMPa cleavage could occur at the inferred site, see Discussion). In both diagrams, cell line differences in the O-glycosite profile are highlighted with a black dot (•). Abbreviations: SP, signal peptide; TM, transmembrane domain; T-166, a known O-glycosite of rhG-CSF produced in CHO cells^17^, A-C, mucin domains; D1-D4, fibronectin domains; S/T, serine-threonine rich region; CHO, Chinese hamster ovary cell line; HEK 293, human embryonic kidney cell line; NS0, mouse myeloma cell line.

The predominant O-glycan form detected on rhG-CSF glycopeptides was a di-sialylated core 1 structure (Table 3). Other observed glycan species were asialylated and mono-sialylated core 1, core 1 structures containing glycolyl neuraminic acid (Neu5Gc), and sialylated core 2 O-glycans. Since O-glycoprofiling using IMPa O-glycoprotease permits detection of intact O-glycans, the presence of post-glycosylation modifications (PGM’s) such as acetylation and sulfation was also assessed. To do this, spectra were searched using an expanded set of O-glycan structures containing sulfate or acetylated sialic acid. rhG-CSF glycopeptides containing acetylated and sulfated O-glycans were identified (Figure S19).

**Table 3.**
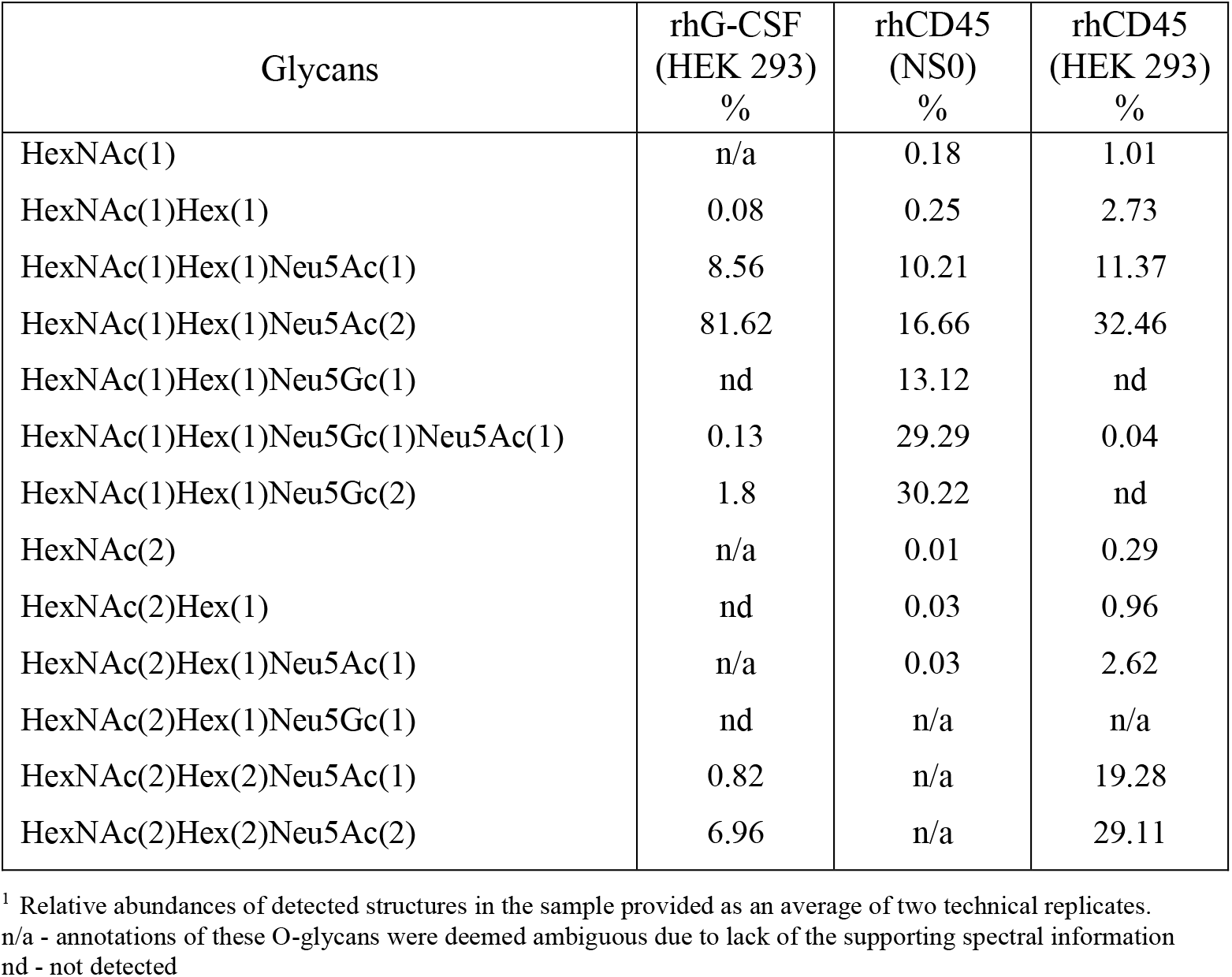
O-Glycan structures detected on rhG-CSF and rhCD45^1^.

| Glycans | rhG-CSF<br>(HEK 293)<br>% | rhCD45<br>(NS0)<br>% | rhCD45<br>(HEK 293)<br>% |
| --- | --- | --- | --- |
| HexNAc(1) | n/a | 0.18 | 1.01 |
| HexNAc(1)Hex(1) | 0.08 | 0.25 | 2.73 |
| HexNAc(1)Hex(1)Neu5Ac(1) | 8.56 | 10.21 | 11.37 |
| HexNAc(1)Hex(1)Neu5Ac(2) | 81.62 | 16.66 | 32.46 |
| HexNAc(1)Hex(1)Neu5Gc(1) | nd | 13.12 | nd |
| HexNAc(1)Hex(1)Neu5Gc(1)Neu5Ac(1) | 0.13 | 29.29 | 0.04 |
| HexNAc(1)Hex(1)Neu5Gc(2) | 1.8 | 30.22 | nd |
| HexNAc(2) | n/a | 0.01 | 0.29 |
| HexNAc(2)Hex(1) | nd | 0.03 | 0.96 |
| HexNAc(2)Hex(1)Neu5Ac(1) | n/a | 0.03 | 2.62 |
| HexNAc(2)Hex(1)Neu5Gc(1) | nd | n/a | n/a |
| HexNAc(2)Hex(2)Neu5Ac(1) | 0.82 | n/a | 19.28 |
| HexNAc(2)Hex(2)Neu5Ac(2) | 6.96 | n/a | 29.11 |

<sup>1</sup> Relative abundances of detected structures in the sample provided as an average of two technical replicates. n/a - annotations of these O-glycans were deemed ambiguous due to lack of the supporting spectral information nd - not detected

### Site-specific O-glycoprofiling of recombinant human CD45

Next, LC-MS/MS analysis of the heavily O-glycosylated CD45 protein was performed. Human CD45 is member C of the class 1 (receptor-like) protein tyrosine phosphatase family (PTPRC). It is a type I transmembrane protein containing a large and highly N- and O-glycosylated extracellular domain. Eight CD45 isoforms can be generated through alternative splicing of the CD45 gene. The isoforms affect the protein’s extracellular domain and thus influence the degree protein O-glycosylation.^18^

We compared the O-glycoprofiles of recombinant human CD45 proteins (rhCD45, extracellular domain (26-575aa) of CD45RABC) expressed and purified from two different cell lines, mouse myeloma cells (NS0) and human embryonic kidney (HEK 293) cells. Each native glycoprotein was digested with IMPa, analyzed by LC-MS/MS and the data searched as described above.

Digestion of rhCD45 expressed in NS0 cells yielded 4614 peptide spectrum matches (PSM’s) and 106 unique glycopeptides. A total of 564 glycoforms of these peptides were identified (Table S3). For rhCD45 from HEK 293 cells, 1455 PSM’s, 94 unique glycopeptides and 533 glycoforms of these peptides were identified. Over 95% of the IMPa-generated rhCD45 peptides in both samples were O-glycopeptides, supporting the notion that these O-glycopeptides could be analyzed without further enrichment or processing (Table S3).

A total of 31 O-glycosites were identified after IMPa digestion of rhCD45 from NS0 cells, versus 38 sites for rhCD45 from HEK 293 cells (Figure 4 and Table S3). Additionally, 11 sites in rhCD45 (NS0) and 8 sites in rhCD45 (HEK 293) were inferred from the protease digestion pattern, but no peptides starting at the prospective sites were observed (Figure 4 and Figure S20). Most of the identified O-glycosites were highly clustered at the N-terminal distal end of the CD45 ectodomain (27-180 aa). This corresponds to a known heavily O-glycosylated region encoded by exons 4, 5 and 6.^18^

The major O-glycan structures identified on rhCD45 (NS0) were core 1 structures containing glycolyl neuraminic acid (Neu5Gc) (Table 3). This is consistent with prior observations that murine myeloma cell lines (NS0, Sp2/0) produce Neu5Gc at significantly higher levels compared to other non-human cell lines like CHO or BHK-21.^29–31^ Other observed species were Tn-antigen, asialylated and mono-, di-sialylated core 1 containing Neu5Ac. Structures that could be extended core 1 or core 2 (HexNAc(2)Hex(1)NeuAc(1), HexNAc(2)Hex(2)NeuAc(1), HexNAc(2)Hex(2)NeuAc(2)) were also annotated by Byonic software on the rhCD45 (NS0) glycopeptides. However, these annotations were deemed ambiguous due to lack of supporting spectral information.

Major O-glycoforms detected on rhCD45 (HEK 293) were mono- and di-sialylated core 1 and core 2 (Table 3). Other observed species were Tn-antigen and asialylated core 1. Very few glycan structures containing Neu5Gc were observed. Finally, O-glycans containing sulfated HexNAc were also identified (Figure S21). Thus, differences in O-glycosite occupancy and notable differences in the O-glycan structures observed were identifiable on the same protein expressed in two different mammalian cell lines.

Considered together our experiments with G-CSF and rhCD45 illustrate that IMPa can be used in a one-step analytical workflow to both identify O-glycosites and the range of intact glycan structures present at each site in purified individual O-glycoproteins, including biotherapeutics. Technical considerations for analyzing complex samples are further addressed in the Discussion section.

## Discussion

In this study, the substrate specificity of the immunomodulating metalloprotease from *Pseudomonas aeruginosa* (IMPa) was interrogated using different peptides and their glycoforms. The presented data show that IMPa can recognize and specifically cleave peptides and proteins carrying O-glycans of different complexity nearly irrespective of the glycan’s surrounding amino acid sequence. We highlight examples of utilization of IMPa in O-glycoprofiling of individual glycoproteins without the need for enrichment of generated glycopeptides. We show that IMPa can be used to identify both O-glycosites and the range of complex O-glycan structures present at each site of a purified glycoprotein.

The current study helps to further the understanding of IMPa function. It has been recently proposed that the N-terminal portion of IMPa contains a domain that specifically recognizes proline at P1 followed by O-glycosylated Ser or Thr.^13^ The structural evidence supporting this model was obtained from crystallization of a catalytically inactive IMPa mutant complexed with an O-glycopeptide. The peptide was bound to the N-terminal domain but failed to occupy the active site of the enzyme. Thus, it was unclear if a substrate’s peptide sequence affects the enzyme’s catalytic activity. Our glycopeptide cleavage data clearly show that IMPa has a broad tolerance for amino acids at P1, indicating that this site is likely not a strict specificity determinant. However, in some cases, the P1 residue (*e*.*g*., Asp, Arg or Ile) may influence catalytic efficiency. Examples from our LC-MS/MS data also support this notion. For example, inefficient cleavage was observed for rhCD45 (HEK293) at Ser146, a position preceded by Ile145 (Table S3). Thus, while digestion with IMPa enables qualitative mapping of O-glycosites and interrogation of O-glycan structures at each site, quantitative analyses of O-glycosite occupancy may be affected by varying IMPa cleavage efficiency at sites preceded by Asp, Arg, or Ile. These aspects of IMPa cleavage specificity must be considered during data interpretation.

Our study also demonstrates how IMPa can be used to map O-glycosites and determine the structure of O-glycans at each site without the need for prior removal of sialic acids. In an analytical workflow, the tolerance of IMPa for a broad range of O-glycan structures generates a diverse range of glycopeptide complexity. In highly complex O-glycoproteins (*e*.*g*., rhCD45 in this study) generated glycopeptides may possess one or more glycosites, and each glycosite may harbor a range of possible glycan structures. Additionally, in highly glycosylated O-glycoproteins, O-glycans are often clustered, a physical feature that is difficult to deconvolute. Several observations in the analysis of rhCD45 helped us better understand IMPa performance in a complex sample. For example, more than half of detected rhCD45 glycopeptides contained multiple putative glycosites and two O-glycans (note: we limited our Byonic searches to a maximum of two O-glycans per peptide). Of these, a significant portion contained two adjacent O-glycosites at the N-terminus of the peptide. This supports the notion that the enzyme does not cleave between two occupied adjacent glycosites (Figure 2A). Other observed multi-glycan peptides had two glycosites in close proximity (e.g., 1-2 amino acids apart) (Figure S20). These peptides suggest that IMPa is likely unable to accommodate substrates with O-glycans on serine or threonine residues in positions P1-P3 relative to a cleavage site. Together, these two peptide classes accounted for most of the detected rhCD45 glycopeptides that contained two glycans per peptide (Table S3). While comprehensive digestion of the O-glycan clusters can be useful, it can also lead to generation of very short glycopeptides that may escape LC-MS analysis or may be difficult to map. Although not specifically addressed in this study, we feel partial proteolysis by IMPa in the clustered regions of O-glycosites (as in rhCD45) may be beneficial for refining O-glycosite profiling of heavily O-glycosylated proteins.

In our analysis of rhCH45, many O-glycosites could only be inferred from the IMPa digest profile, with no corresponding glycopeptides being identified. This may occur for several reasons: i) the peptide might ionize poorly, ii) the peptide might fragment poorly, iii) the peptide may be too large or too small, and iv) the peptide may have another post-translational modification (e.g, N-glycan) that was not accounted for in databases searches. In analysis of rhCD45, inferred O-glycosites map to Ser70, Thr72, Thr73 and Thr74 (Figure S20B). Corresponding glycopeptides containing these glycosites were not identified in our standard searches. Interestingly, there are three known N-glycosylation sites (Asn80, Asn92, Asn97) located in proximity to these sites. As N-glycans were not removed from rhCD45 proteins before IMPa digestion and N-glycan structures were not included in our Byonic searches due to their complexity, it is plausible that peptides containing both N- and O-glycans were missed in our analysis.

For peptides bearing more than one Ser or Thr, deducing the O-glycosite position and O-glycan structure on each glycopeptide can be difficult using only HCD fragmentation. Fragmentation of glycopeptides using collision-based techniques (CID/HCD) often result in losses of glycans on peptide backbone fragments, hampering unambiguous O-glycosite assignment. Detailed knowledge of IMPa specificity can aid in data interpretation. Its highly specific cleavage at the N-terminus of a modified Ser/Thr and its inability to cleave between two adjacent or closely situated O-glycosites provides valuable guidance for accurately mapping O-glycosites on each glycopeptide. Additionally, it may help to resolve ambiguities in annotated spectra. For instance, applying IMPa specificity rules may help to annotate O-glycosite(s) on peptides with multiple potential sites (Figure S22). However, for unambiguous assignment of two or more specific O-glycan structures at each glycosite HCD fragmentation data may not be enough, for which fragmentation using electron transfer dissociation methods (ETD or EThcD) may be needed. These methods can preserve glycan modifications and can generate relevant fragmentation data.^32–35^ We envision that interrogation of O-glycoproteins by digestion with IMPa, coupled with LC-MS/MS analysis using EThcD dissociation would provide superior information for characterization of O-glycopeptides.

## Acknowledgements

We wish to honor the memory of NEB founder and scientist Dr. Donald Comb who was devoted to basic research at NEB, through which this work was made possible.

## Author contributions statement

S.V. conceived the study; S.V. and C.H.T designed experiments; S.V., H.G., E.M., X.S. performed experiments; S.V. and H.G. analyzed data; C.M. and C.R. performed mass spectrometry analyses; S.V. and C.H.T wrote the manuscript. All authors reviewed and approved the final version of the manuscript.

## Supporting Information

Supporting information accompanies this paper: additional experimental details and methods, chromatographic profiles of the analyzed peptides/glycopeptides, representative MS/MS spectra, and datasets of the identified glycopeptides.

