## Supporting Methods, Table S1 and Figures for "A broad-specificity O-glycoprotease that enables improved analysis of glycoproteins and glycopeptides containing intact complex O-glycans"

New England Biolabs, 240 County Road, Ipswich, MA 01938, USA

### Supporting Methods

#### Information S1. Preparation of recombinant O-endoglycoproteases

*Cloning of BT4244 and StcE O-endoglycoproteases*

Codon optimized DNA fragments encoding the BT4244 protease from *Bacterioides thetaiotaomicron*'s (GenBank Accession Q89ZX7) and StcE protease from *Escherichia coli* O157:H7 (GenBank BAA31757.3) were synthetized by Integrated DNA Technologies (Coralville, IA). All primers and plasmids were designed using the NEBuilder® Assembly Tool (http://nebuilder.neb.com/). The DNA fragments encoding a truncated version of BT4244 (274 to 857 aa) containing a C-terminal $His_6$-tag with terminal vector-specific overlapping sequences and StcE (36-898 aa) containing a N-terminal $His_6$-tag with terminal vector-specific overlapping sequences were amplified by PCR. Amplified DNA fragments were purified by gel extraction and cloned into the vector pJS119k using the NEBuilder HiFi DNA Assembly cloning Kit (New England Biolabs, Ipswich, MA). Competent *E. coli* NEB 10-β cells (New England Biolabs) were transformed with 2 µL of the reaction mix. The resulting pJS119k/BT4244, pJS119k/StcE plasmids were purified using the Monarch Plasmid Miniprep Kit (New England Biolabs) and sequenced.

*Expression and purification of recombinant BT4244 and StcE proteases*

For protein expression, an overnight cultures of *E. coli* NEB Express cells carrying pJS119k/BT4244 or pJS119k/StcE were diluted 1:100 in 2 L of LB medium supplemented with 40 µg/mL kanamycin and grown to 0.4 $OD_{600}$ at 30°C. The expression of recombinant protein was induced by addition of isopropyl-β-thiogalactopyranoside (IPTG) to a final concentration of 0.4 mM and shaking for 4 h at 30°C. The cells were harvested by centrifugation and suspended in 80 ml of 20 mM Tris-HCl, pH 8.0 containing 500 mM NaCl. Cells were disrupted using a high shear homogenizer at 18,000 psi (PL300, Dyhydromatics, Maynard, MA). The total extract was clarified by centrifugation for 1 hour at 27,000 x *g*. The cleared lysate was loaded onto a $Ni^{2+}$ immobilized by chelation with nitrilotriacetic acid (NTA) metal affinity chromatography column (HisTrap FF 5 ml, GE Healthcare, Chicago, IL), washed with 50 column volumes (CV) of 10 mM imidazole, and eluted with a 20 CV gradient consisting of 20 mM Tris-HCl, pH 8.0 containing 500 mM NaCl and 10 to 175 mM imidazole. Pooled elution fractions containing pure protein were dialyzed against 20 mM Tris-HCl, pH 8.0 and concentrated using Vivaspin 20 30,000 Da MWCO concentrators (Sartorius Stedim Biotech). Purified proteins were stored at -20°C.

**Information S2. Preparation of recombinant hGALNT1**

*Plasmid and strain construction*

All primers and plasmid were designed using the NEBuilder® Assembly Tool (http://nebuilder.neb.com/). The construct encoding 29-559 aa of the human Polypeptide N-Acetylgalactosaminyltransferase 1 (GALNT1) containing C-terminal His-tag was amplified from human cDNA using primers 5'-cgagaaaagagaggccgaagctagtgaatgcaacaaatgtgatg and 5'-gagcggccgccccttcaacctcagtgatgatgatgatgatggaatatttctggcagggtg-3'. pD912 vector backbone was amplified using primers 5'-agcttcggcctctcttttctcgagag-3' and 5'-ggttgaaggggcggccgctca-3'. DNA fragments were purified using the QIAquick Gel Extraction Kit (Qiagen, Valencia, CA). To assemble the expression vector, 25 ng of pD912 backbone and 100 ng of DNA fragment to be inserted were added to an NEBuilder® HiFi DNA Assembly Master Mix reaction (New England Biolabs, Ipswich, MA). Two microliters of the assembly mix were used as DNA template to amplify a linear expression cassette using primers 5'-gctcattccaattccttctattag-3' and 5'-gagctccaatcaagcccaataac-3'. The amplified linear expression cassette was purified by gel extraction. The resulting secretion construct encoded hGALNT1 with an N-terminal yeast α-MF signal sequence and a C-terminal HIS-tag under control of the constitutive GAP promoter. For expression in yeast, the amplified linear expression cassette (0.1 μg) was introduced into *Pichia pastoris* (*Komagataella phaffii*) Mut$^S$ competent cells by electroporation. Transformants were selected by growth on YPD agar plates supplemented with 500 mg/L zeocin for 3-4 days at 30°C.

*Expression and purification of rhGALNT1*

*P. pastoris* cultures were grown in BMGY medium consisting of 10 g/L yeast extract, 20 g/L peptone, 13.4 g/L YNB, 0.4 mg/L D-biotin, 0.1 M potassium phosphate buffer pH 6 with 1% (w/v) glycerol as a carbon source. The yeast cells were grown for 48 h at 30°C with shaking. Cell culture supernatant was harvested by centrifugation, concentrated and buffer exchanged to 20 mM Tris-HCl, pH8.0 containing 0.3M NaCl using Vivaspin 20 30 kDa filters. The resulting sample was loaded onto a $Ni^{2+}$ immobilized by chelation with nitrilotriacetic acid (NTA) metal affinity chromatography column (HisTrap FF 5 ml, GE Healthcare, Chicago, IL), washed with 50 column volumes (CV) of 10 mM imidazole, and eluted with a 20 CV gradient consisting of 20 mM Tris-HCl, pH 8.0 containing 0.3 M NaCl and 10 to 175 mM imidazole. Pooled elution fractions containing pure protein were dialyzed against 20 mM Tris-HCl, pH 8.0 and concentrated using Vivaspin 20 30 kDa MWCO concentrators (Sartorius Stedim Biotech, Göttingen, Germany). Purified protein was stored at -20°C.

### Supporting Tables

**Table S1.** A set of 13 common mammalian O-glycans used for Byonic searches

| Glycan | Mass |
|---|---|
| HexNAc(1) | 203.0794 |
| HexNAc(1)Hex(1) | 365.1322 |
| HexNAc(2) | 406.1587 |
| HexNAc(2)Hex(1) | 568.2116 |
| HexNAc(1)Hex(1)NeuAc(1) | 656.2276 |
| HexNAc(1)Hex(1)NeuGc(1) | 672.2225 |
| HexNAc(2)Hex(1)NeuAc(1) | 859.3070 |
| HexNAc(2)Hex(1)NeuGc(1) | 875.3019 |
| HexNAc(1)Hex(1)NeuAc(2) | 947.3230 |
| HexNAc(1)Hex(1)NeuGc(1)NeuAc(1) | 963.3179 |
| HexNAc(1)Hex(1)NeuGc(2) | 979.3129 |
| HexNAc(2)Hex(2)NeuAc(1) | 1021.3598 |
| HexNAc(2)Hex(2)NeuAc(2) | 1312.4552 |

**Table S2. (Excel file).** Identification of site-specific O-linked glycopeptides from rhG-CSF(HEK 293).

**Table S3. (Excel file).** Identification of site-specific O-linked glycopeptides from rhCD45(NS0) and rhCD45(HEK 293).

### Supporting Figures

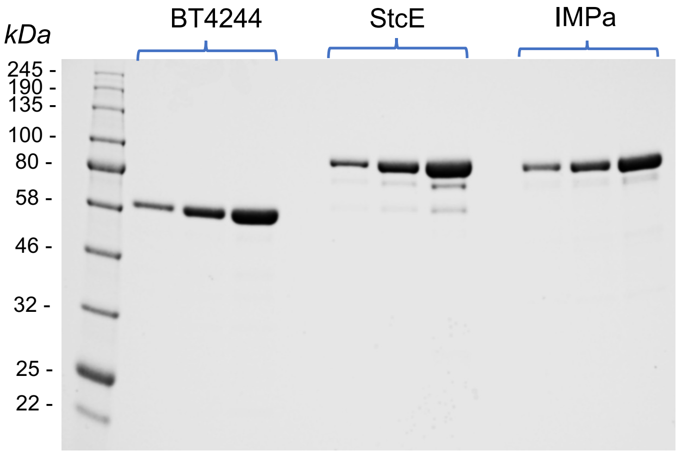

**Figure S1.** Expression and purification of StcE and BT4244. Purified StcE, BT4244 proteins and IMPa (O-Glycoprotease, NEB) were separated by SDS-PAGE and stained with SimplyBlue SafeStain.

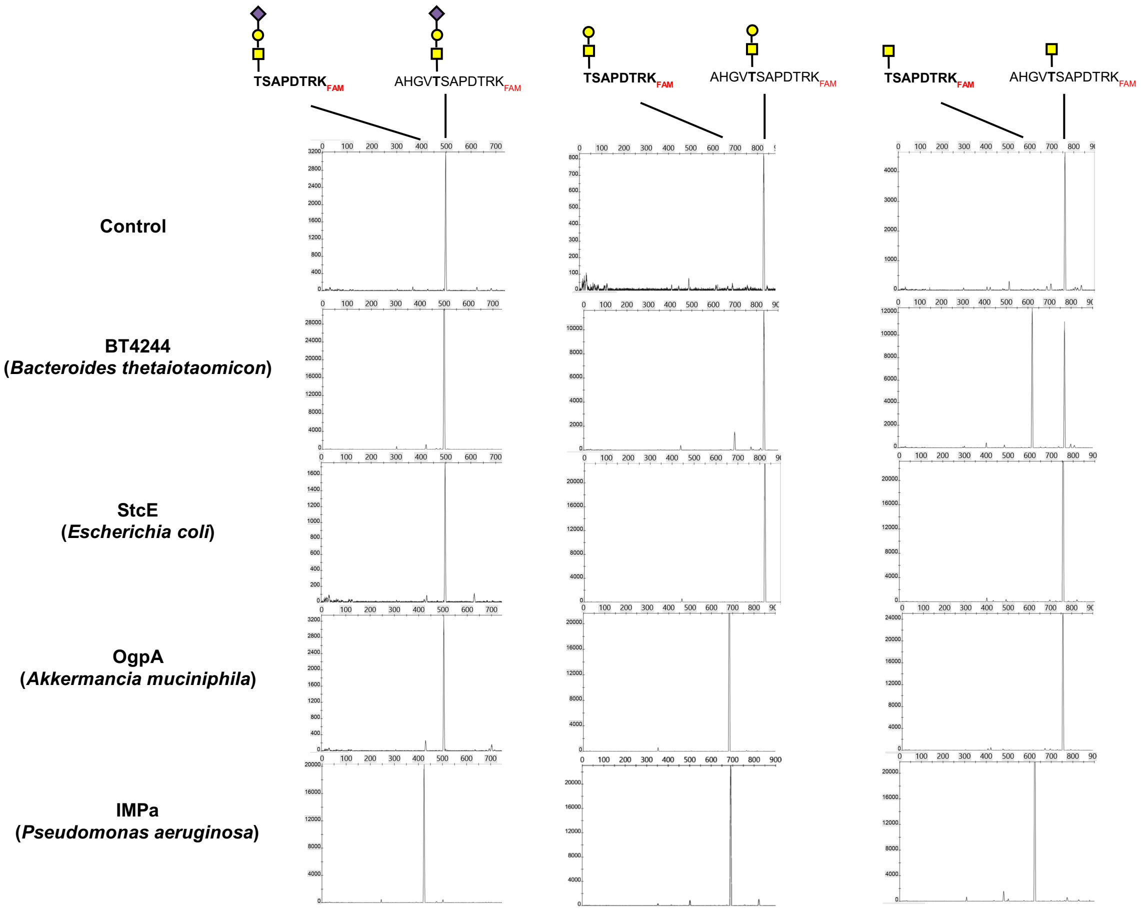

**Figure S2.** Treatment of O-glycosylated synthetic peptides with O-endoproteases. Synthetic peptides containing different O-glycans were incubated with BT274, StcE, OgpA and IMPa enzymes. Reaction mixtures were analyzed by CE.

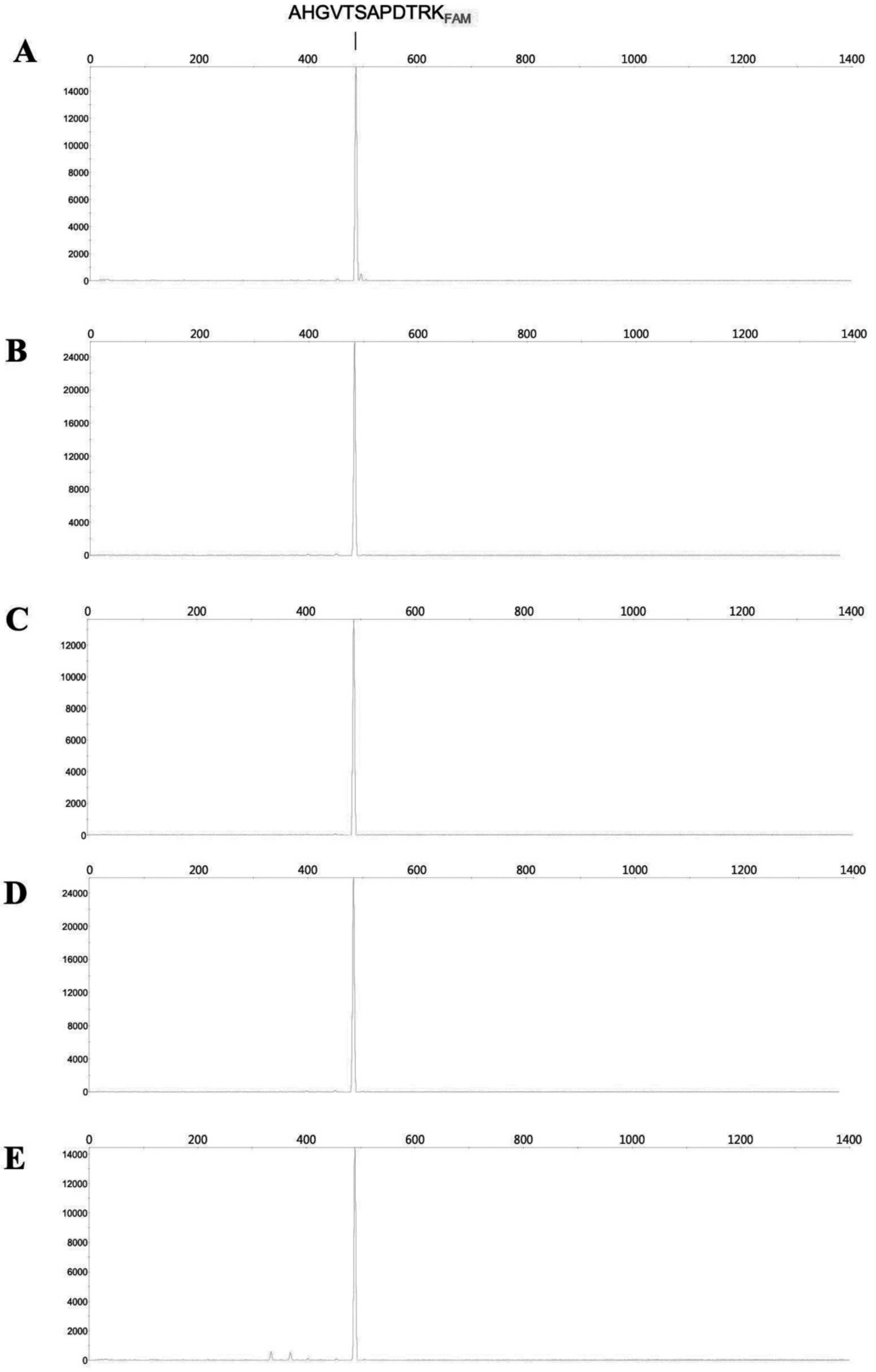

**Figure S3.** Treatment of non-glycosylated synthetic peptide with O-endoproteases. Synthetic peptide (A) was incubated with BT274 (B), IMPa (C), OgpA (D) and StcE (E). Reaction mixtures were analyzed by CE.

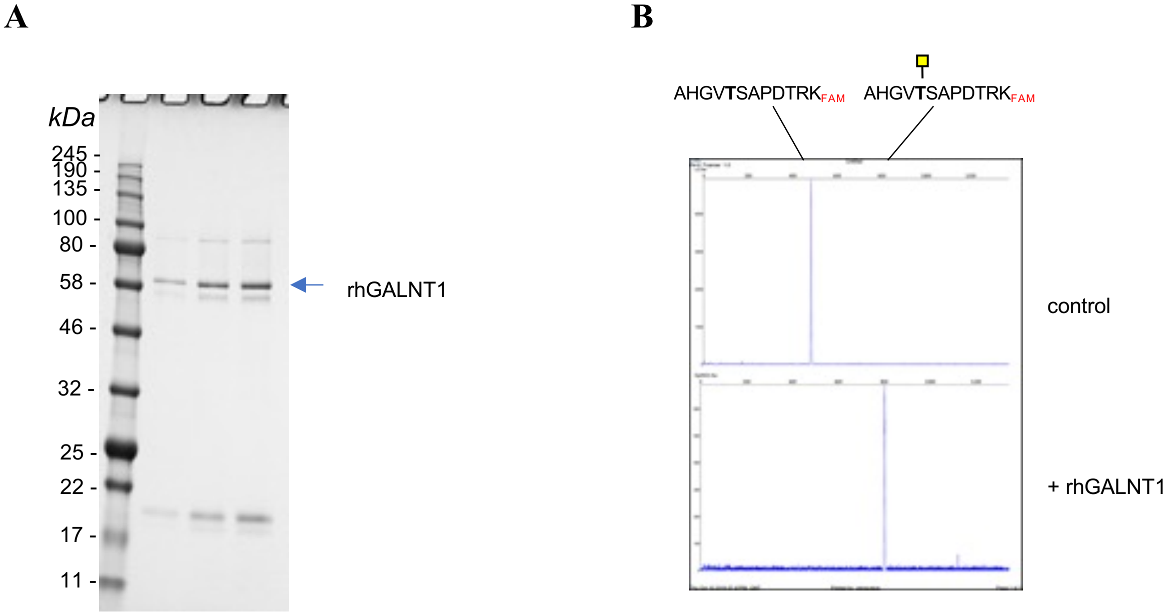

**Figure S4.** Expression and purification of rhGALNT1. Soluble catalytic domain of human GALNT1 containing C-terminal His-tag was expressed in *Pichia pastoris* and purified using Ni-NTA resin. Purified rhGALNT1 was analyzed by SDS-PAGE and stained with SimplyBlue SafeStain (A) or used in the activity assay with FAM-labeled peptide and UDP-GalNAc. Reactions were analyzed by CE (B).

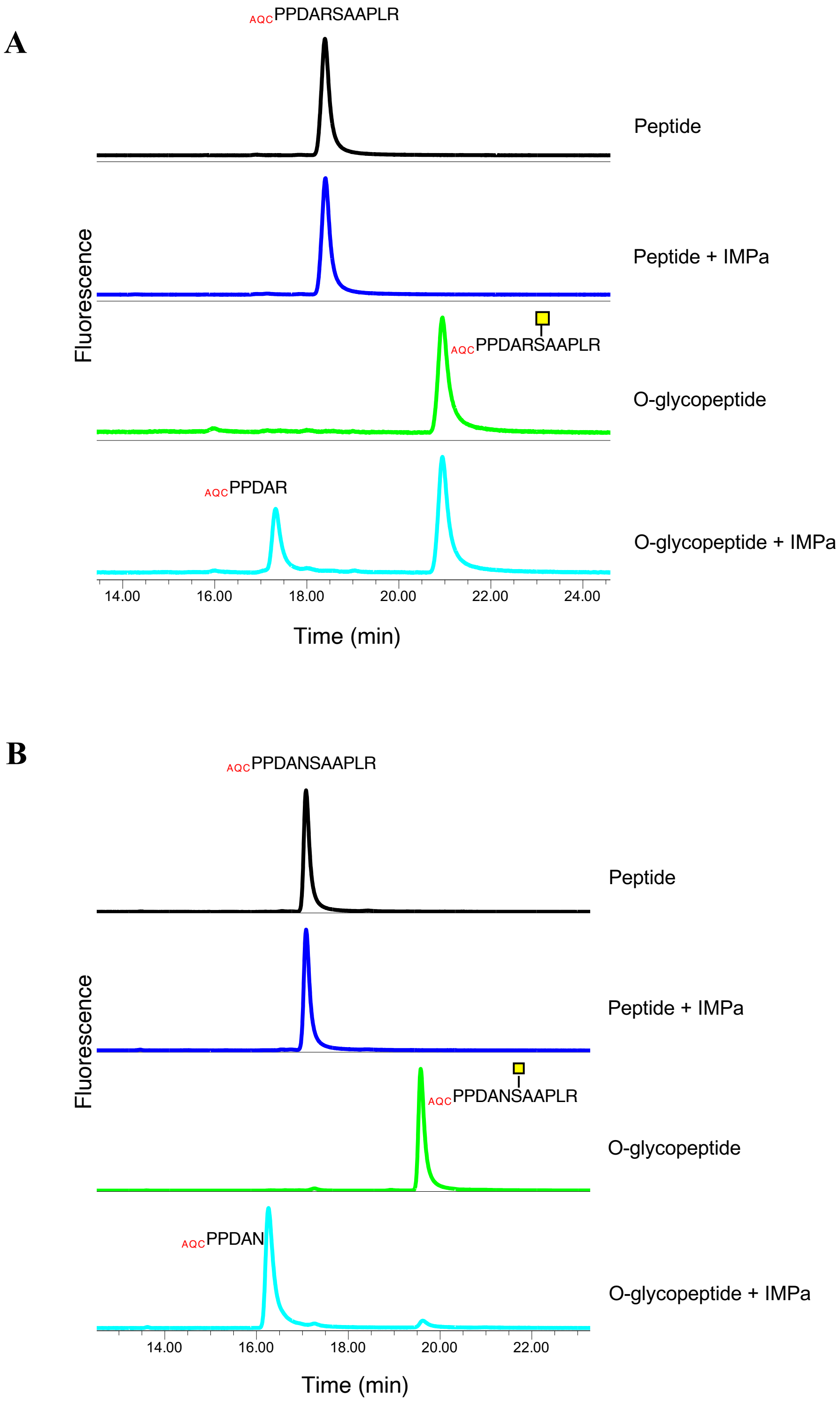

**Figure S5.** IMPa digests of the EPO peptide variants containing a different amino acid at P1 position. (A) R at P1, (B) N at P1.

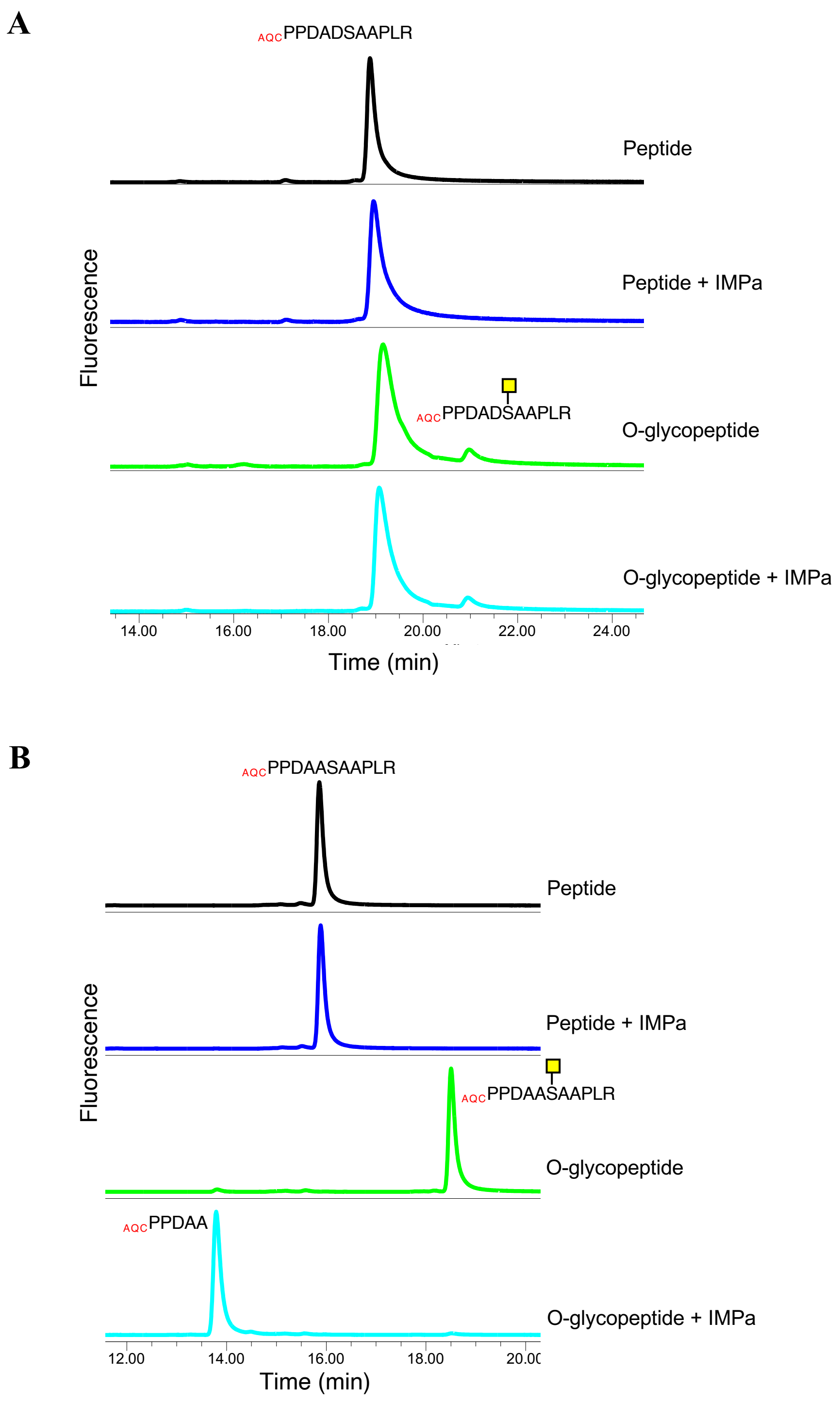

**Figure S6.** IMPa digests of the EPO peptide variants containing a different amino acid at P1 position. (A) D at P1, (B) A at P1.

A

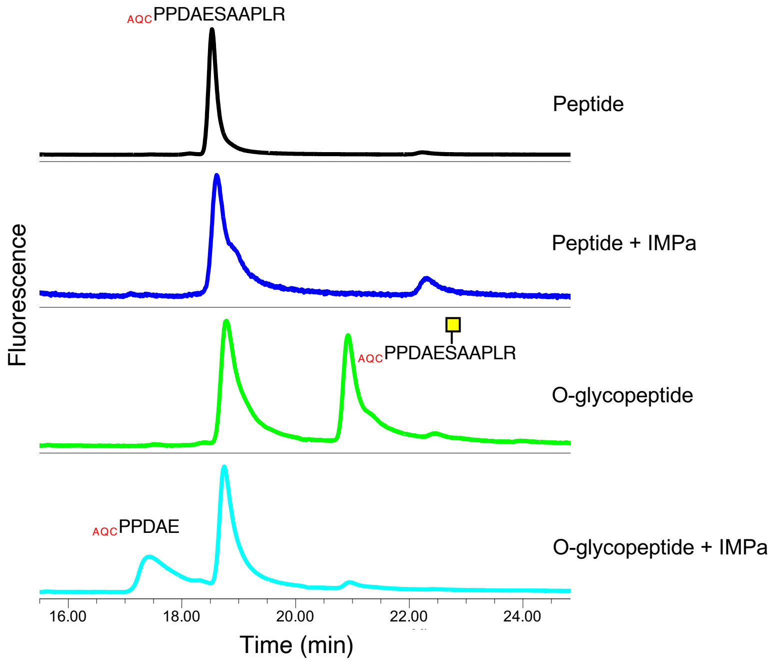

B

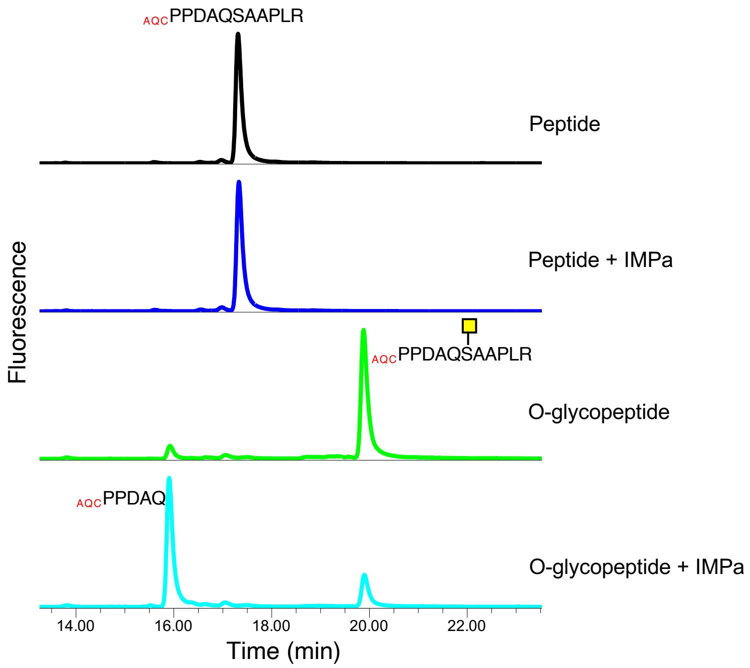

**Figure S7.** IMPa digests of the EPO peptide variants containing a different amino acid at P1 position. (A) E at P1, (B) Q at P1.

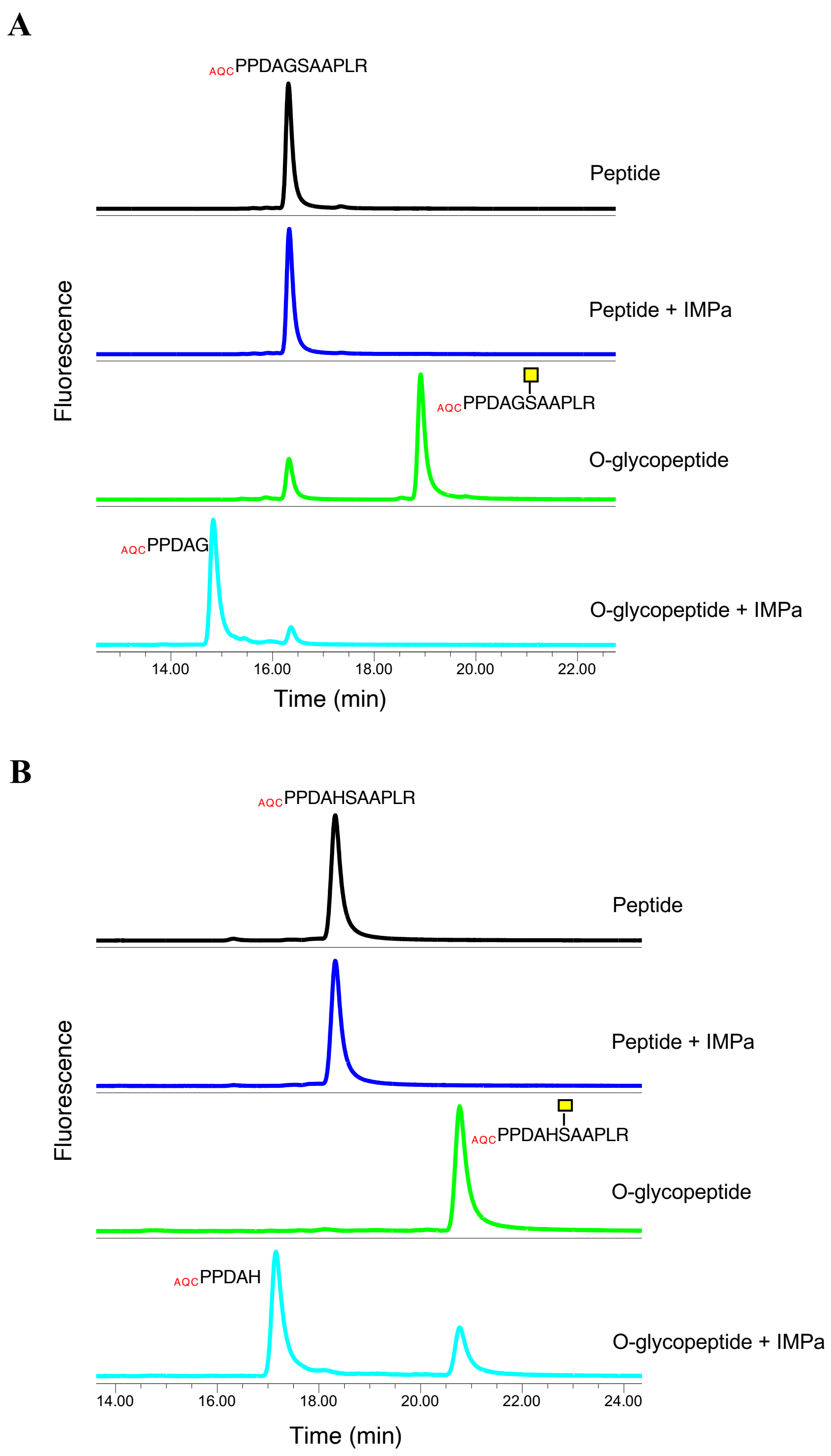

**Figure S8.** IMPa digests of the EPO peptide variants containing a different amino acid at P1 position. (A) G at P1, (B) H at P1.

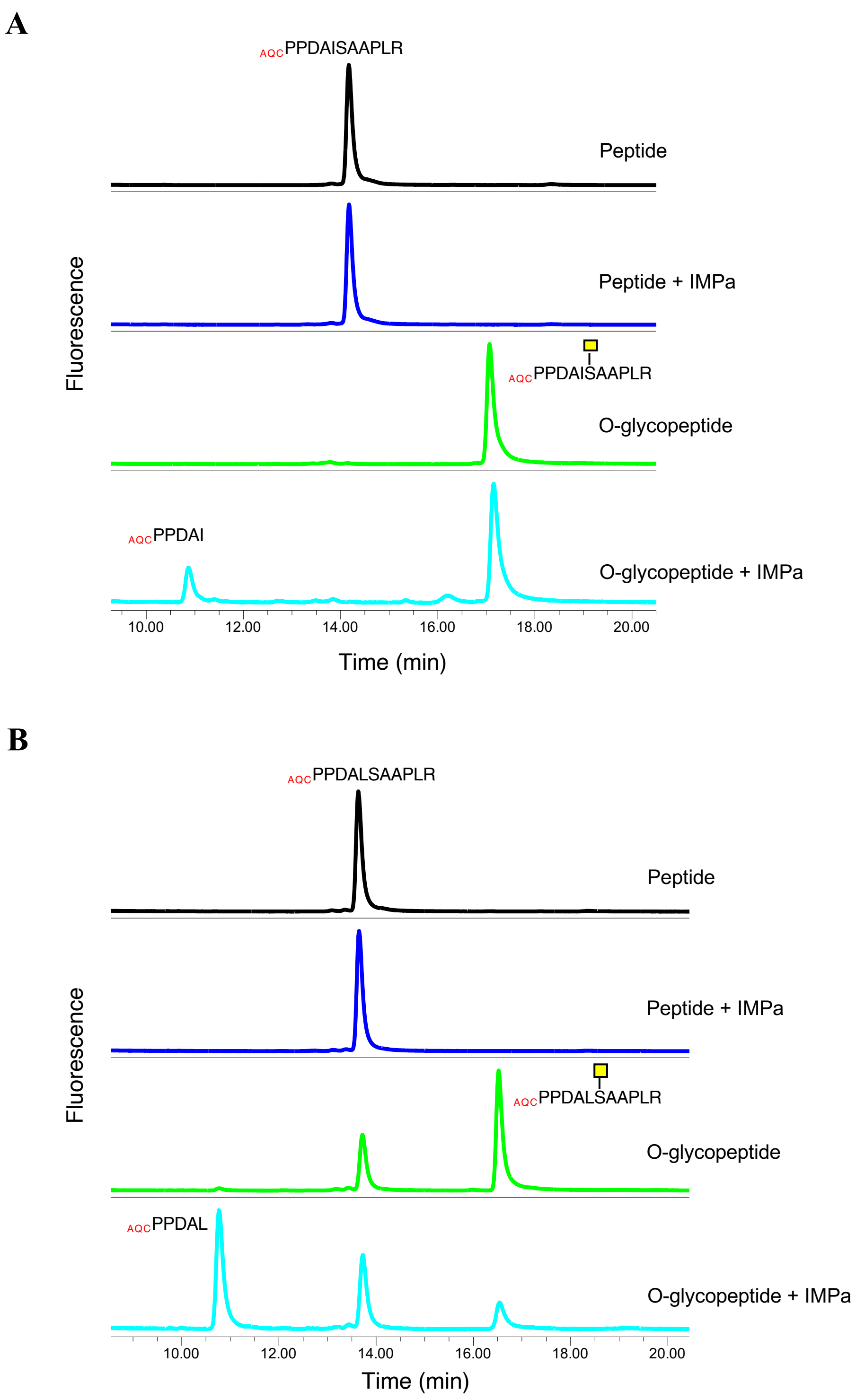

**Figure S9.** IMPa digests of the EPO peptide variants containing a different amino acid at P1 position. (A) I at P1, (B) L at P1.

A

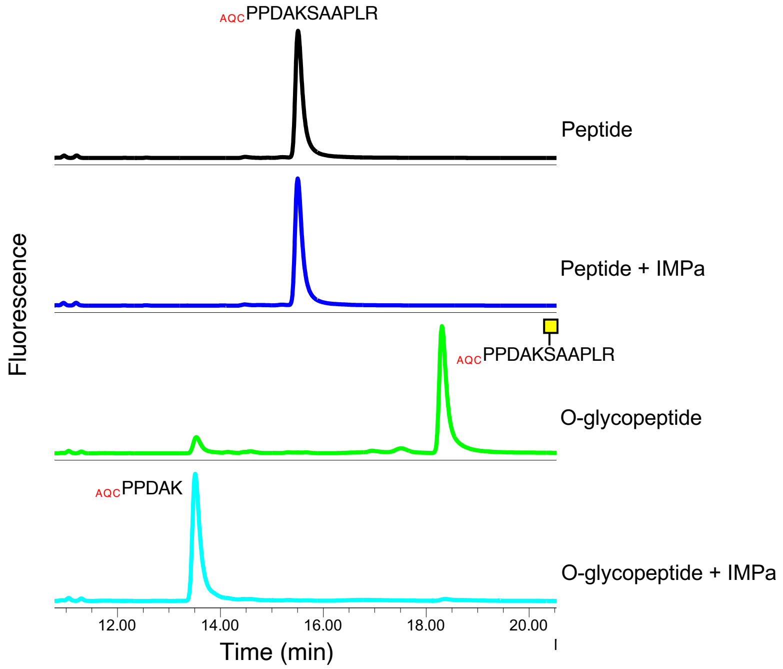

B

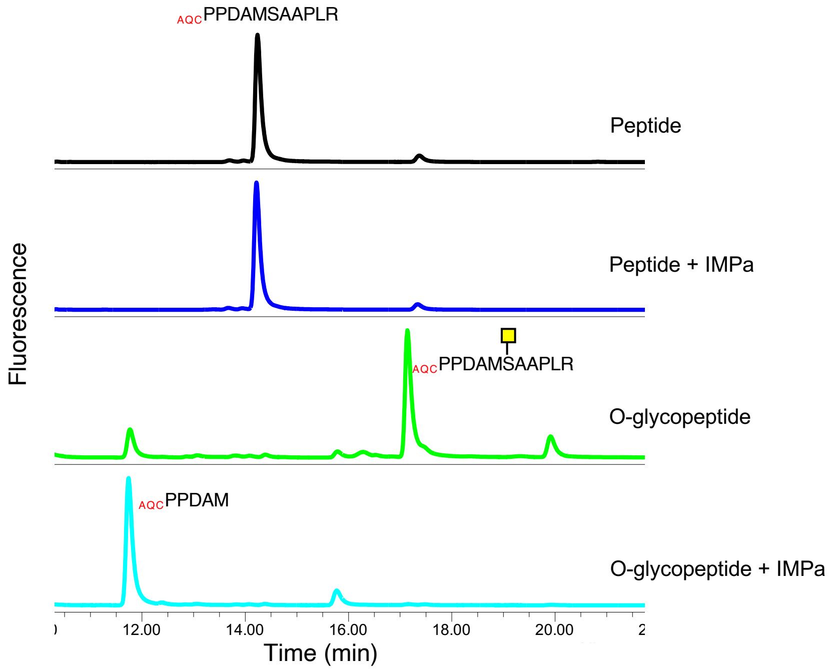

**Figure S10.** IMPa digests of the EPO peptide variants containing a different amino acid at P1 position. (A) K at P1, (B) M at P1.

A

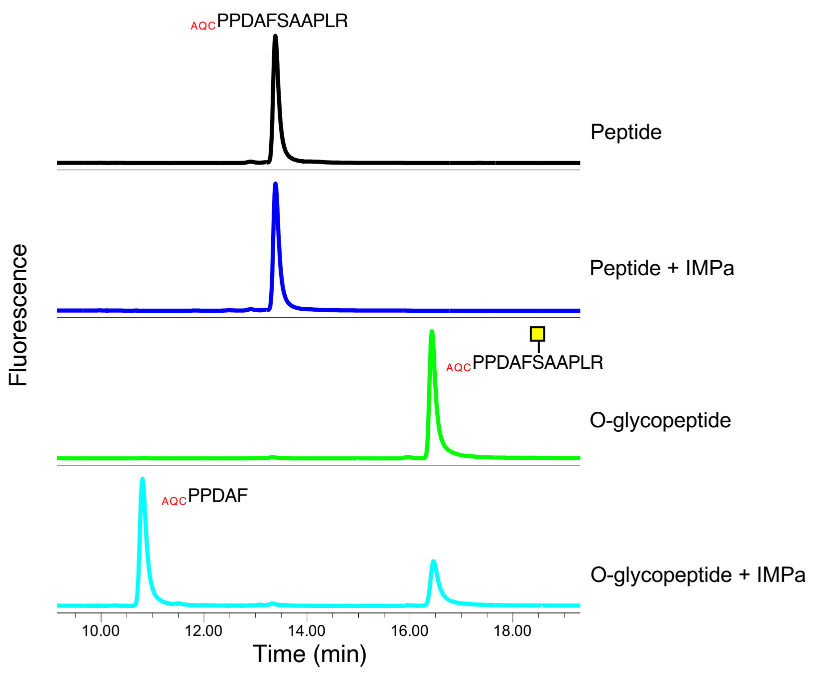

B

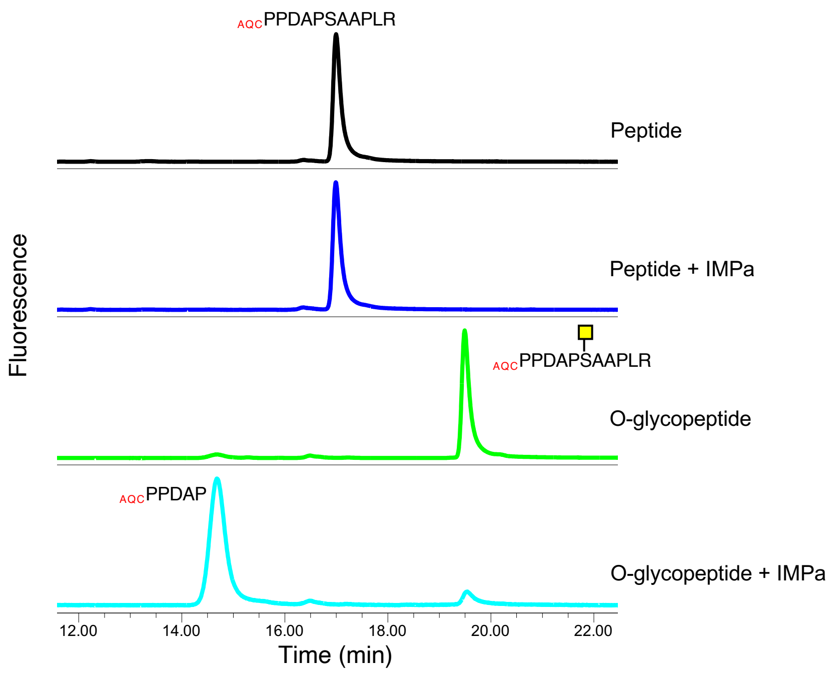

**Figure S11.** IMPa digests of the EPO peptide variants containing a different amino acid at P1 position. (A) F at P1, (B) P at P1.

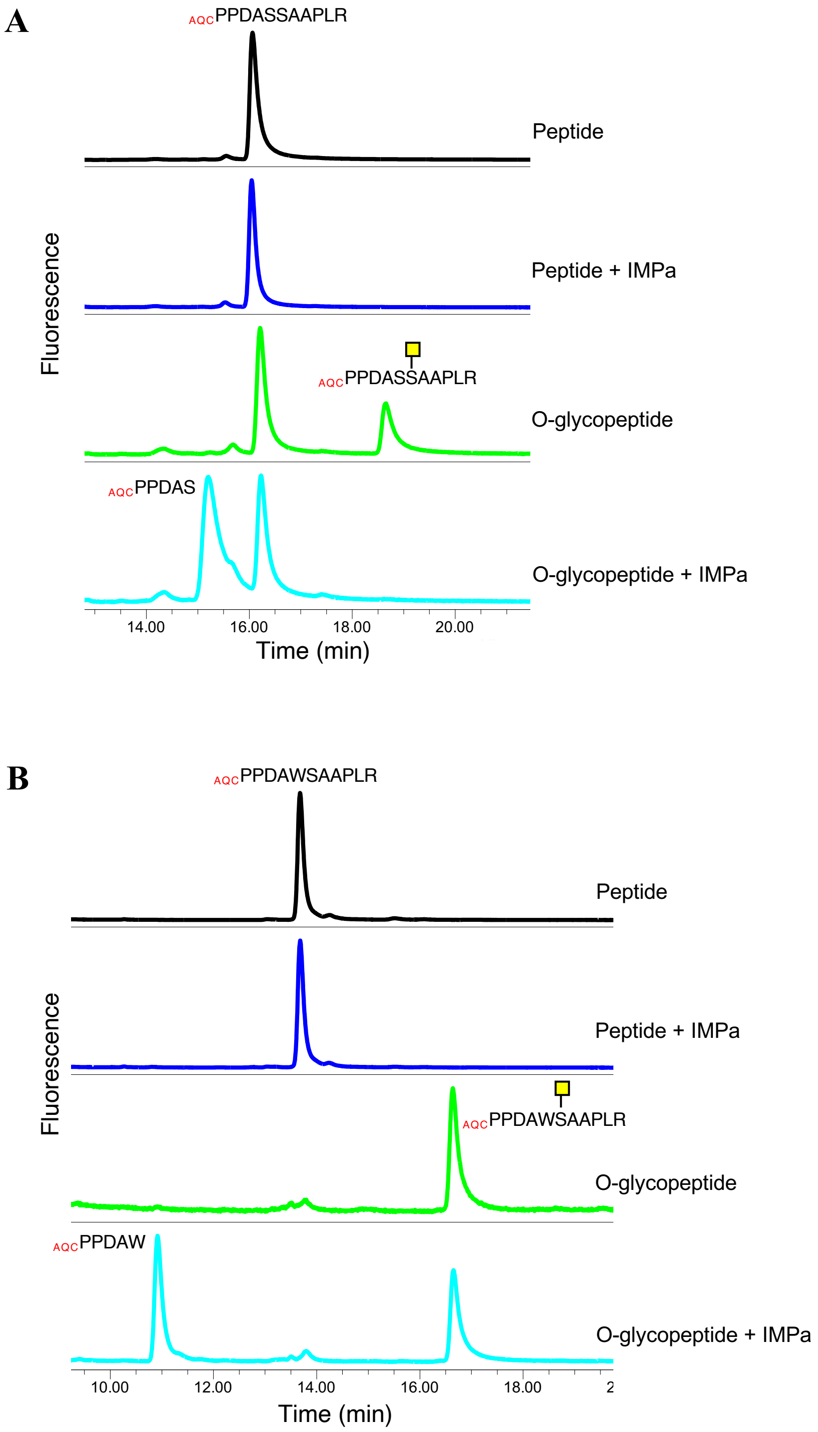

**Figure S12.** IMPa digests of the EPO peptide variants containing a different amino acid at P1 position. (A) S at P1, (B) W at P1.

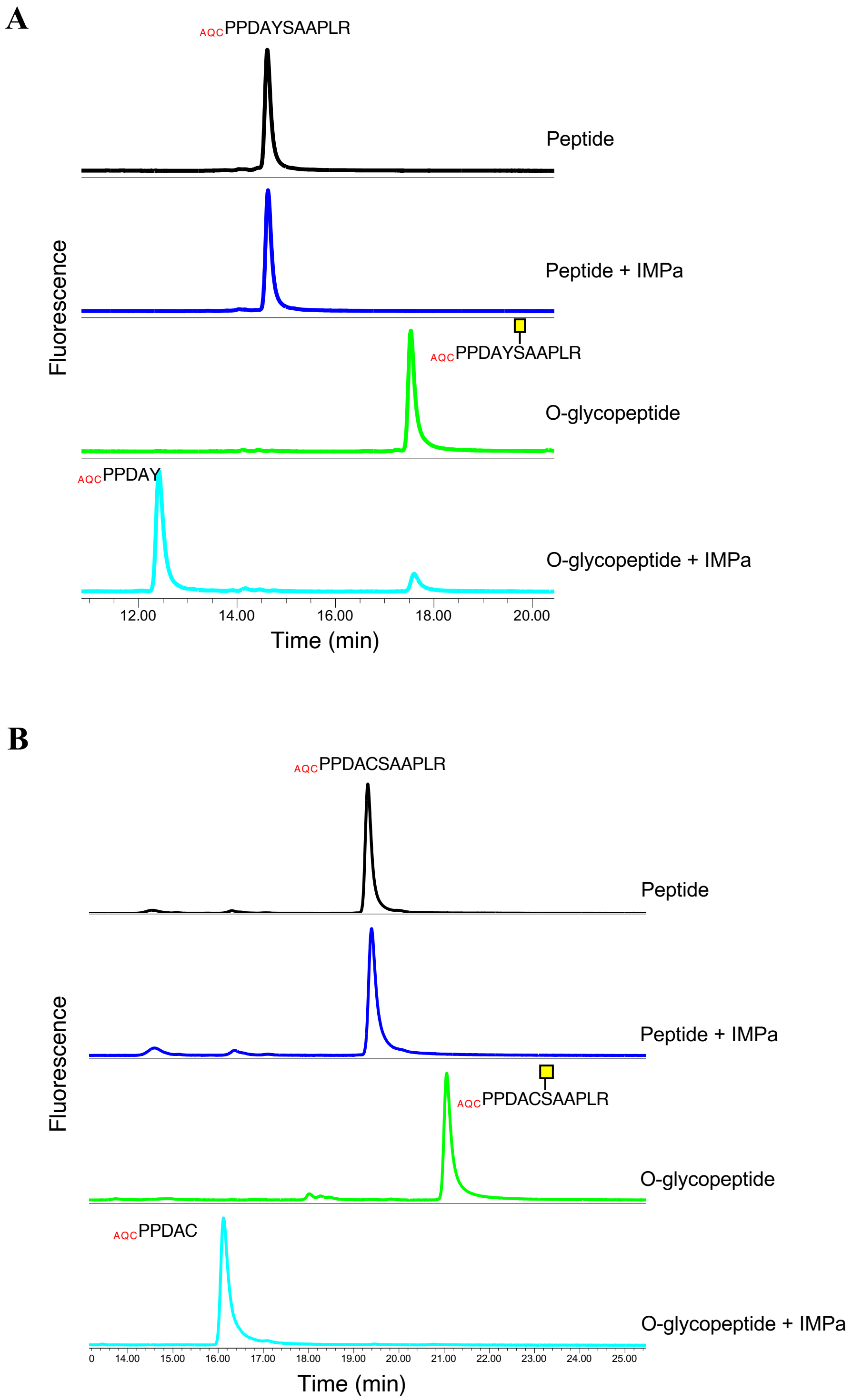

**Figure S13.** IMPa digests of the EPO peptide variants containing a different amino acid at P1 position. (A) Y at P1, (B) C at P1.

**A**

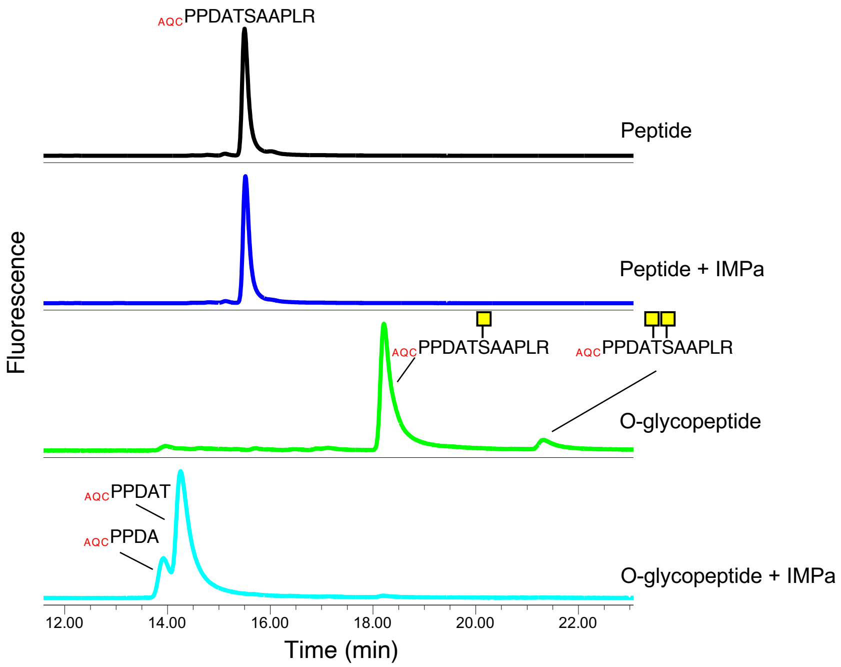

**B**

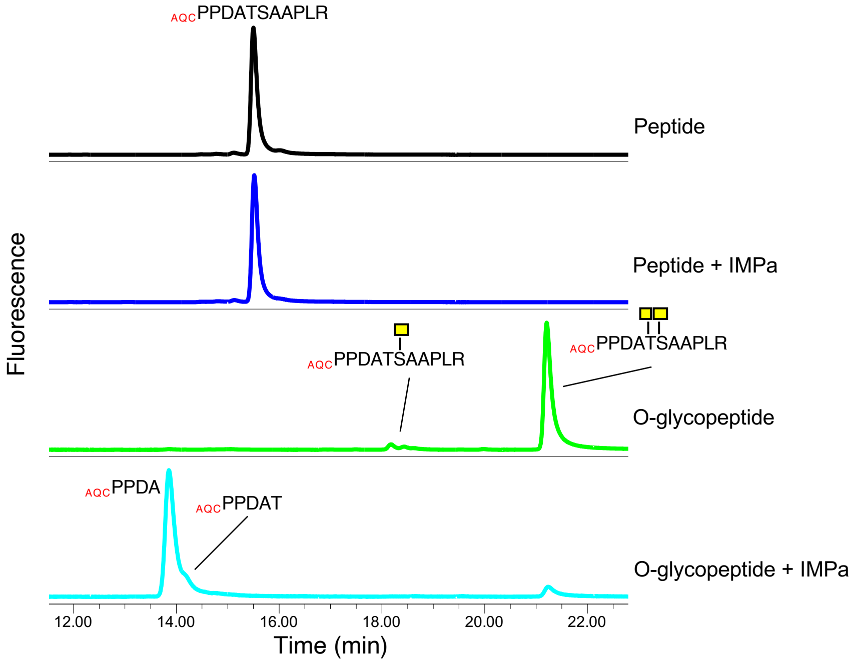

**Figure S14.** IMPa digests of the EPO peptide variants containing a different amino acid at P1 position. (A) T at P1; the peptide incubated with rhGALNT1 for 4h, (B) T at P1; the peptide incubated with rhGALNT1 for 24h.

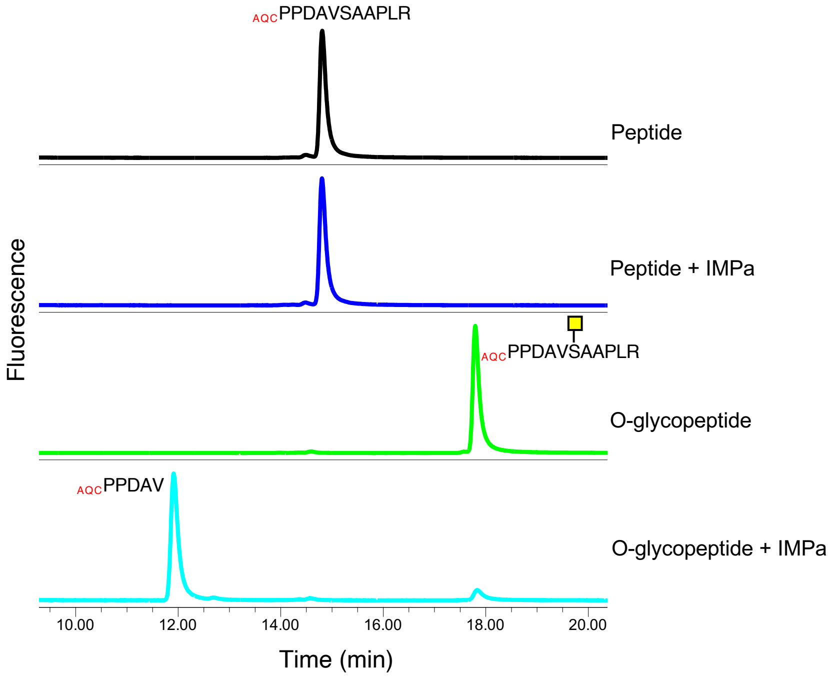

**Figure S15.** IMPa digests of the EPO peptide variant that contains V at P1 position.

A

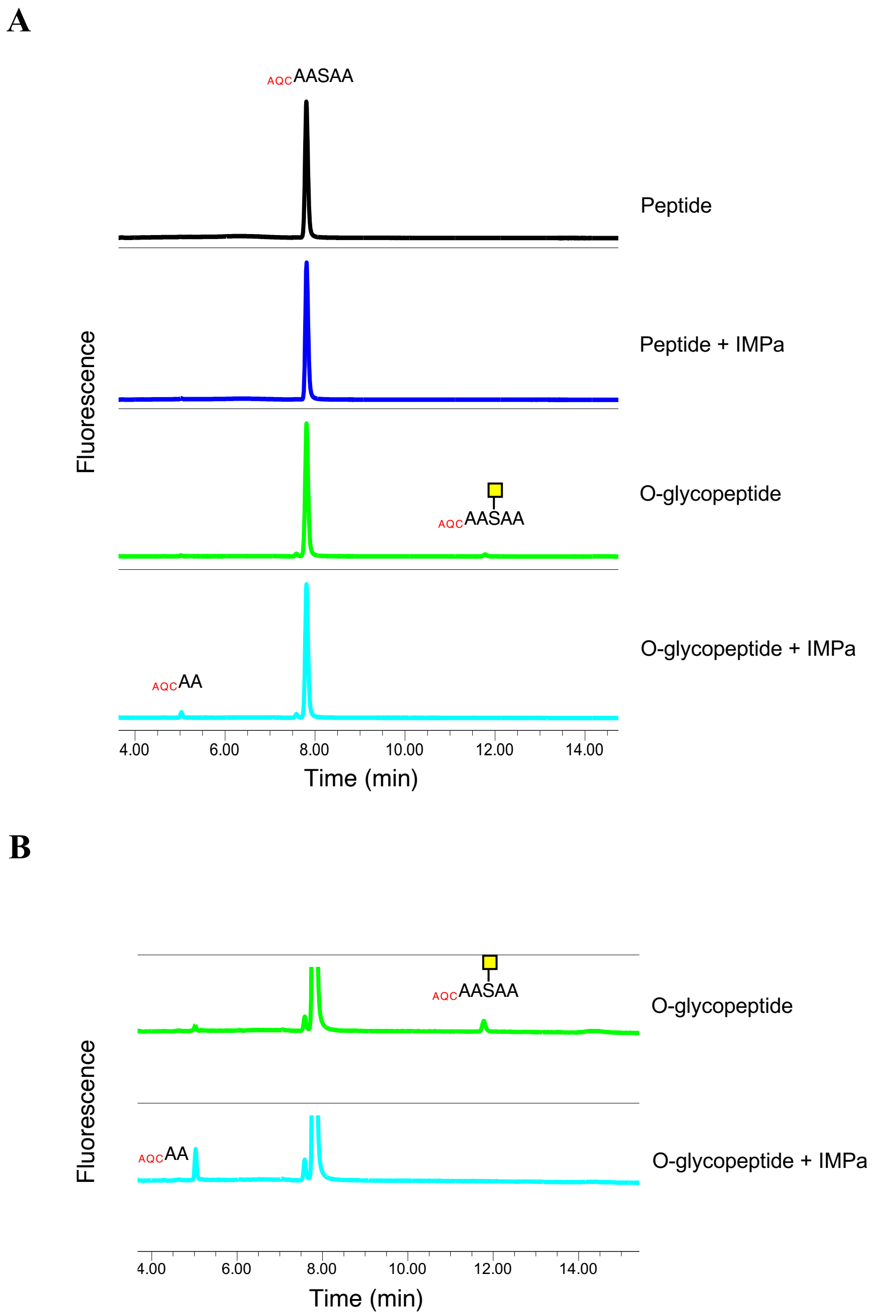

**Figure S16.** IMPa digest of an EPO peptide variant containing 2 amino acid residues next to the cleavage site. (A) Raw chromatogram, (B) Zoomed chromatogram.

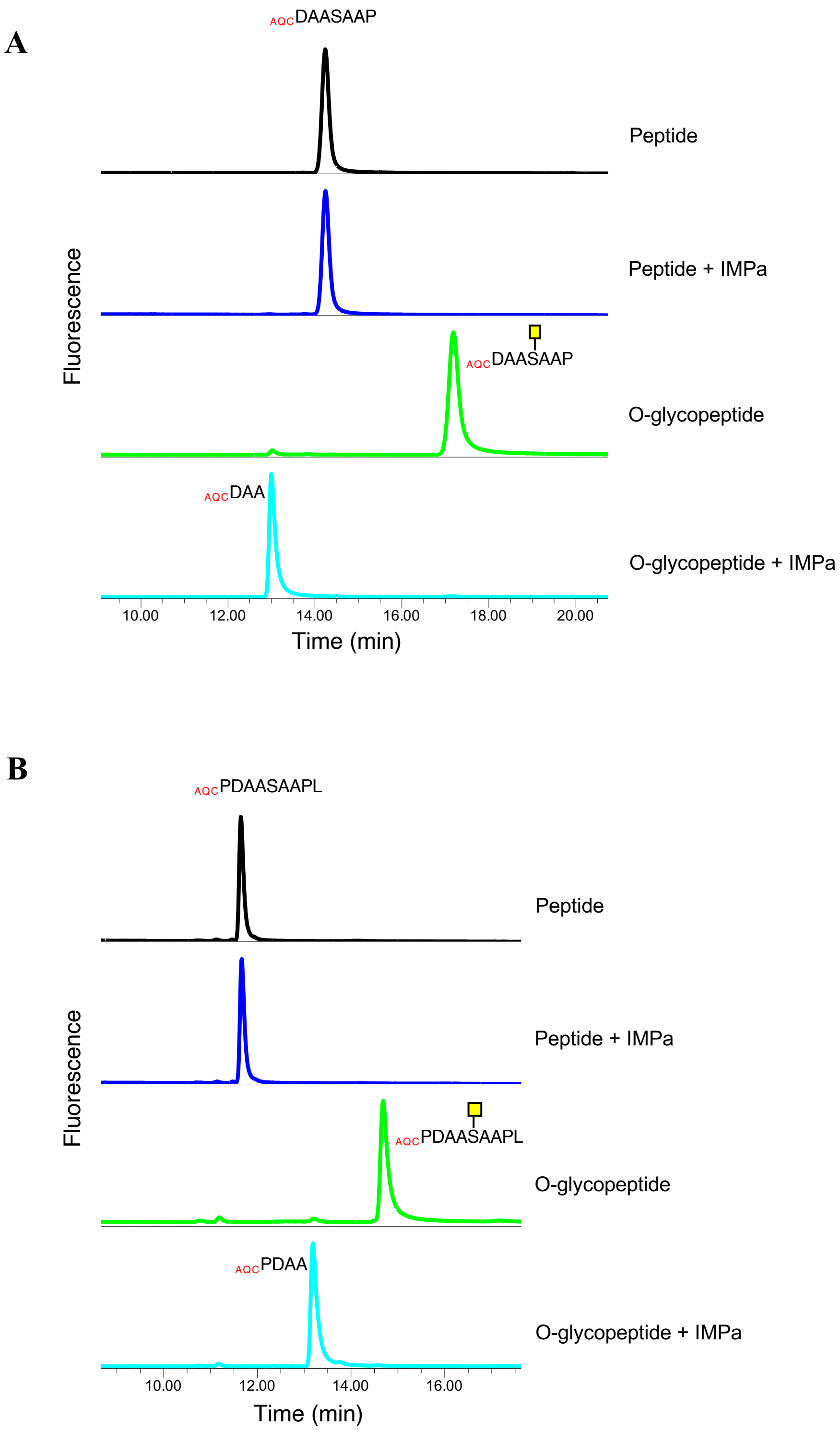

**Figure S17.** IMPa digests of EPO peptide variants containing (A) 3 or (B) 4 amino acid residues next to the cleavage site.

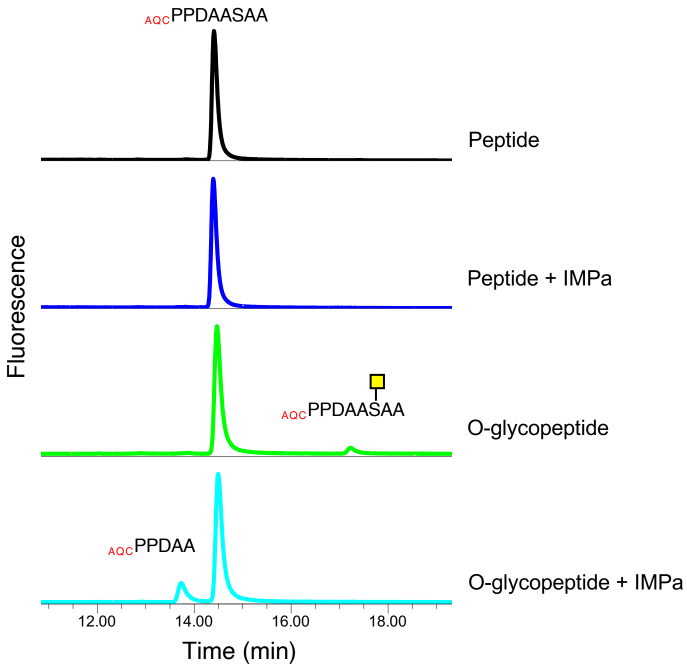

**Figure S18.** IMPa digest of an EPO peptide variant containing 5 amino acid residues upstream of the cleavage site and 2 amino acid residues – downstream of the cleavage site.

A

B

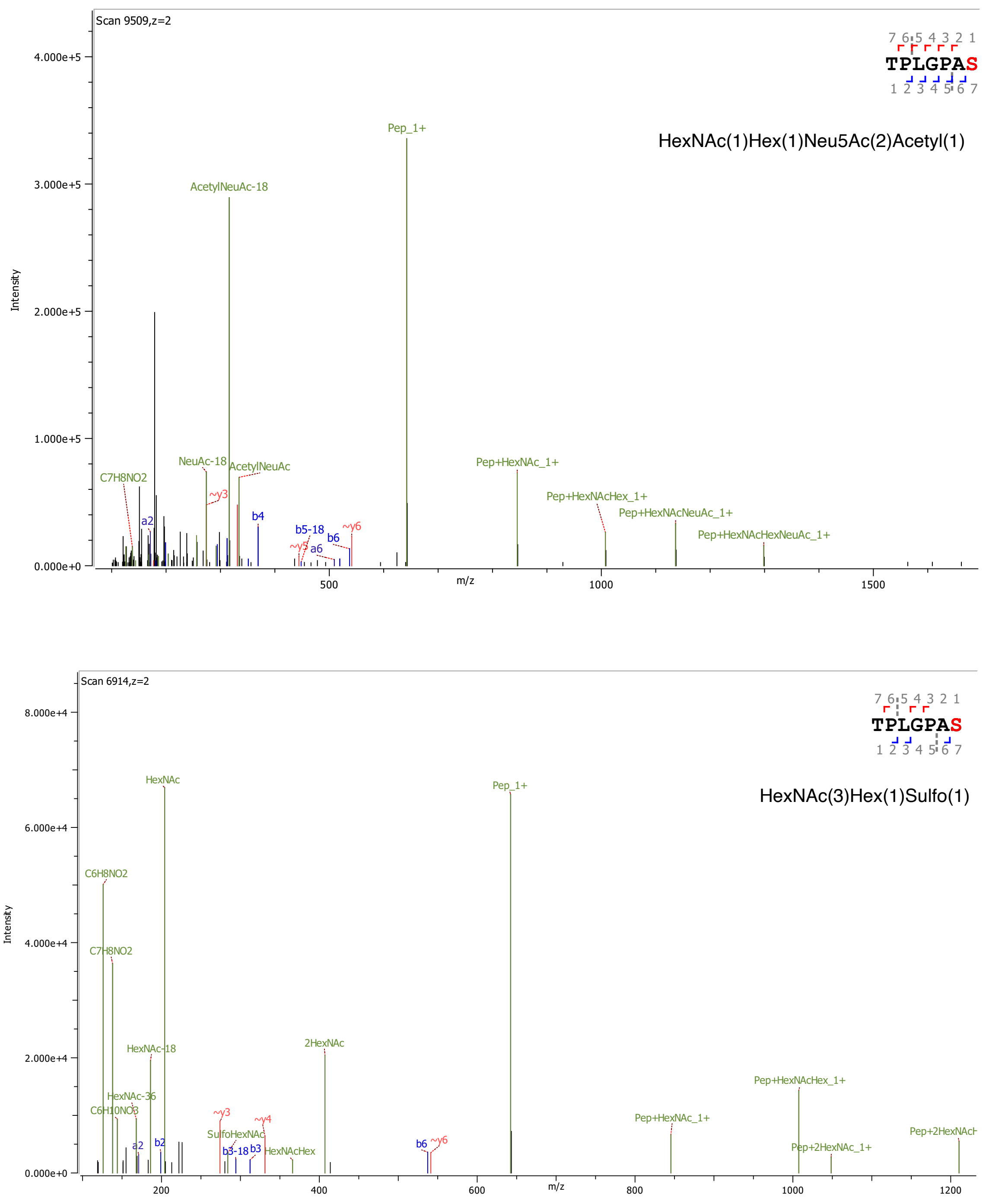

**Figure S19.** MS/MS spectra of representative rhG-CSF glycopeptides containing acetylated (A) and sulfated (B) O-glycan structures. In both cases, the software assigned O-glycosites on C-terminal Ser37 (shown in red) with no confidence (due to the insufficient spectral information). The corrected O-glycosite position on IMPa-generated glycopeptides is N-terminal threonine (Thr31).

A

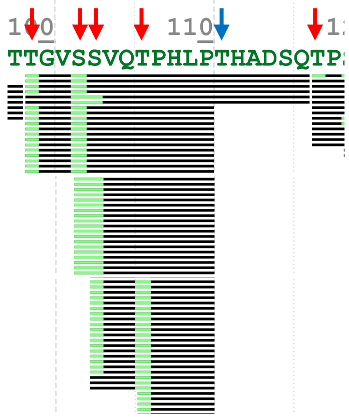

B

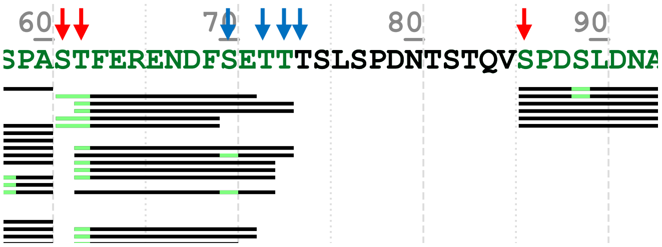

**Figure S20.** Examples of the protein coverage observed in the short rhCD45 regions that contain a cluster of O-glycosites (Byonic view output). Black bars represent detected PSM's (single or group). Byonic annotations of the O-glycosites are shown in green. Arrows indicate location of confirmed (red) and inferred (blue) O-glycosites. (A) Example of the inferred glycosite at Thr111, (B) Example of the multiple inferred O-glycosites located near known N-glycosite (Asn80).

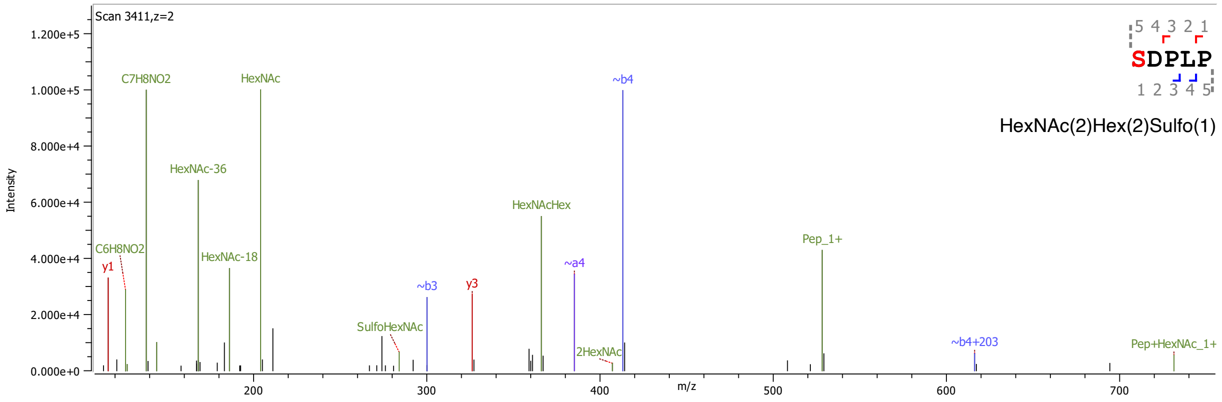

**Figure S21.** MS/MS spectra of the representative CD45 glycopeptide containing sulfated O-glycan structure HexNAc(2)Hex(2)Sulfo(1).

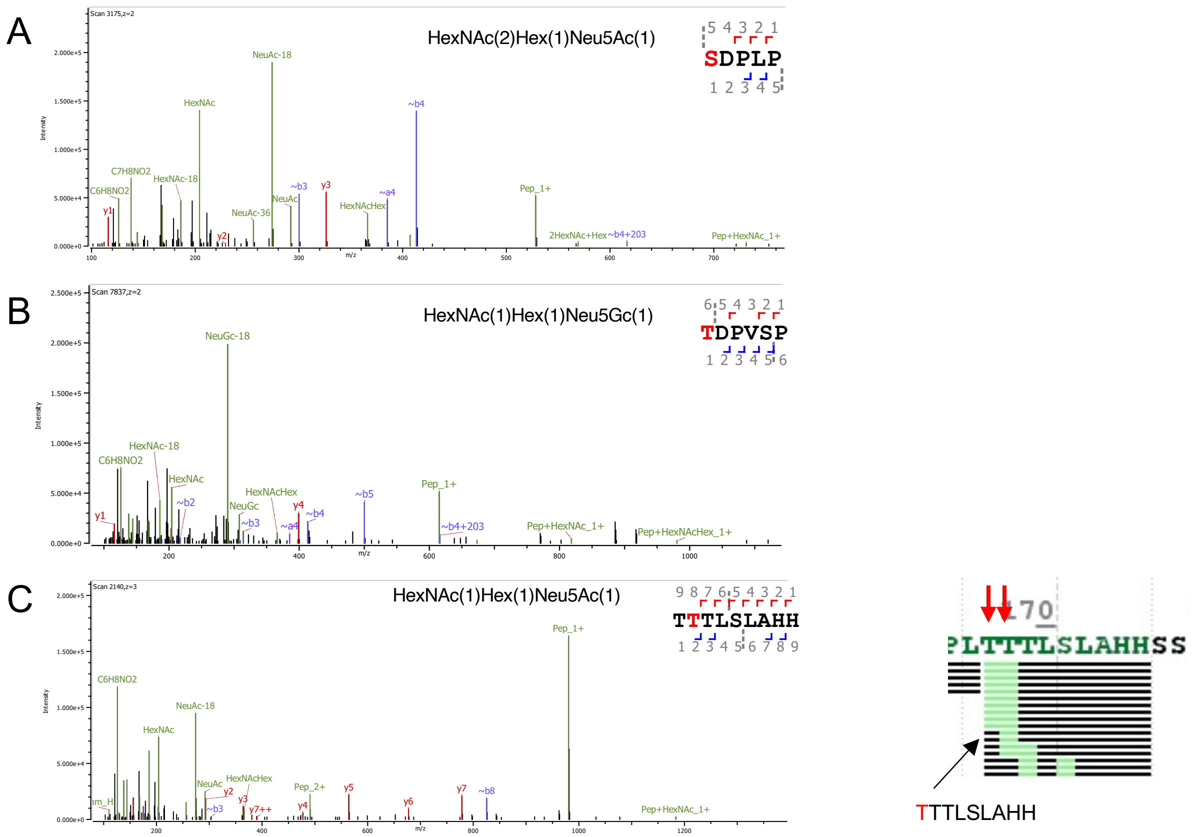

**Figure S22.** Examples of annotations of O-glycosites on the identified rhCD45 glycopeptides (based on available spectral information and known IMPa specificity). Representative spectra are of the glycopeptide containing single O-glycosite (A) and the glycopeptide containing two potential O-glycosites and one O-linked glycan (B). In both cases, the high confidence annotations of N-terminal O-glycosite by Byonic software are based on the observed peptide fragments with partial glycan (in this case, b4 ion with HexNAc (~b4 + 203). These assignments are also supported by IMPa cleavage specificity. (C) Representative spectra of the glycopeptide containing multiple potential O-glycosites and one O-linked glycan. Due to limited spectral information available (no peptide fragments with partial glycan), Byonic assignment of the O-glycosite on the second Thr from N-terminus is highly ambiguous (no confidence). The observed IMPa cleavage pattern and enzyme specificity indicate that the O-glycan is located on the N-terminal Thr residue of this peptide.
